# Assessing the Cognitive Status of Drosophila by the Value-Based Feeding Decision

**DOI:** 10.1101/2020.08.27.267955

**Authors:** Chih-Chieh Yu, Ferng-Chang Chang, Yong-Huei Hong, Jian-Chiuan Li, Po-Lin Chen, Chun-Hong Chen, Tzai-Wen Chiu, Tsai-Te Lu, Yun-Ming Wang, Chih-Fei Kao

## Abstract

Decision making is considered an important aspect of cognitive function. Impaired decision making is a consequence of cognitive decline caused by various physiological conditions, such as aging and neurodegenerative diseases. Here we exploited the value-based feeding decision (VBFD) assay, which is a simple sensory-motor task, to determine the cognitive status of Drosophila. Our results indicated the deterioration of VBFD is notably correlated with aging and neurodegenerative disorders. Restriction of the mushroom body (MB) neuronal activity partly blunted the proper VBFD. Furthermore, using the Drosophila polyQ disease model, we demonstrated the impaired VBFD is ameliorated by the dinitrosyl iron complex (DNIC-1), a novel and steady nitric oxide (NO)-releasing compound. Therefore we propose that the VBFD assay provides a robust assessment of Drosophila cognition and can be used to characterize additional neuroprotective interventions.

## Introduction

Decision making is the act of choosing between available options in facing a need or a problem. The process of decision making usually involves several steps, including the identification of a need and its potential options, evaluation of options, decision making and acting, and finally a review of the decision that may assist the prospective decision making when a similar need/problem is encountered. Decision making is generally considered as a high-level cognitive process (*1, 2*). The most intriguing part of the decision making process is the evaluation of accessible options, which may be based on values, preferences, and past experience of the decision maker. The underlying process and neural circuits of decision making are currently topics of intense study. A common experimental paradigm for elucidating neural decision making in mammals is the two-alternative forced choice task (2AFC). Two alternative options are concurrently presented to the test subject. Within a defined time, the test subject has to choose between two alternatives. Distinct features of the two alternatives and creative experimental designs have been exploited to study the specific behavior dynamics of choice under different physiological conditions and, most importantly, the involved neural elements.

Similar two-choice assays have been adapted by the Drosophila model to study diverse aspects of the feeding behavior (i.e., two-choice feeding assay) (*3–5*). Such studies have allowed scientists to achieve substantial progress in understanding the neuronal and molecular mechanisms that modulate feeding decisions. Particularly, feeding decisions are implemented in the nervous system on multiple levels, from the peripheral chemosensory organs to the central brain (*3, 6–8*). Many neuronal and molecular mechanisms regulating insect feeding decisions have been uncovered (*9–12*). Interestingly, previous studies have demonstrated wild-type (WT) fruit flies are able to differentiate the nutritional values of two sugar solutions and learn to associate an odorant paired with the nutritious sugar solution (*3, 6, 7*).

The advantageous decision for most young WT flies to ingest the metabolizable sugar, but not the non-metabolizable sugar allows the starved flies to quickly regain the nutritional homeostasis. Most importantly, the efficacy of feeding decision based on the caloric contents of two sugar solutions is robust and no pre-conditioning is required. To be able to make the proper feeding decision underscores the neural mechanisms governing the decision making process, which is regarded as a unique process of cognitive function. The efficacy of proper feeding decision may therefore be used to promptly and faithfully evaluate the cognitive status of fruit flies.

In this study, we explored the efficacy of value-based feeding decision (VBFD) in Drosophila under selected physiological conditions. Our results indicated the efficacy of VBFD is susceptible to the chronological ages, alterations of life expectancy, and neurodegenerative disorders. The characterization of simple and complex VBFDs also enabled us to investigate distinct levels of Drosophila cognitive function and identify neural circuits related to the making of VBFD. Lastly, we further demonstrated the VBFD assay is an effective approach to identify compounds that have protective functions to disease-associated cognitive disorders.

## Results

### Deterioration of proper value-based feeding decision (VBFD) in the aged Drosophila

To meet the acute nutritional needs, fast and reliable mechanisms are necessary for fruit flies to evaluate the nutritional values of food substances and determine the subsequent feeding behavior. That is to keep on eating the same food source or switch to another food source with higher nutritional value. Making such VBFD relies on the proper function of central brain mechanisms, which may represent a unique spectrum of cognitive processes. To demonstrate VBFD, the two-choice feeding assay was used to determine the feeding decision of fruit flies (*3–5*). Briefly, in this assay, food deprived fruit flies were presented with two sugar solutions, the metabolizable sucrose and the non-metabolizable arabinose. The relative nutritional values of two sugars were verified by their life supporting capability as the sole source of food (Fig. S1A). Each sugar solution was colored with a blue or a red dye, and the feeding decision was scored by the dye accumulation in the abdomen of individual flies (Fig. 1A). This quantitative measurement of feeding decision indicated the starved 10 day-old w^1118^ adult flies exhibit a strong preference for the metabolizable sucrose (92~97.5% of total tested flies) over the non-metabolizable arabinose (2.5~8%), suggesting young WT flies can efficiently evaluate the caloric contents of sugars and make appropriate VBFD within the short feeding period (Fig. 1B). Moreover, the capability to make proper VBFD was equally effective in both genders (Figs. 1B and S1C). Next, we asked if VBFD remains intact in the aged flies. The survival analysis revealed the median lifespan of w^1118^ flies is ~63 days in females and ~68 days in males (Fig. S1B). We therefore chose two additional time points before the median lifespan to characterize the efficacy of VBFD. Intriguingly, while 40 day-old starved flies recognized and preferred the nutritious sucrose, this capability was significantly deteriorating in 60 day-old flies of both sexes (Figs. 1B and S1C). The population of flies consuming only the non-nutritious arabinose significantly increased to 7~10.5% in 40 day-old flies and to 23~25% in 60 day-old flies. Furthermore, we also noted, as aging progresses, the population of Non-eater flies also increased (from 0% in 10 day-old flies to 11~20% in 60 day-old flies), suggesting this phenotype may be associated with aging. To resolve the acute nutritional needs, food deprived flies have to quickly identify and ingest food substances that are palatable and nutritious. Therefore, the Non-eater flies appeared insensitive to the hunger-driven signals. To explore if the decline of making appropriate VBFD is caused by the inability to sense sugar, we assayed the feeding decision between the sweet sugar solution (sucrose or arabinose) and plain water. Unlike the VBFD between the sucrose and arabinose, the sweet sensation and the preference to either sweet sugar were much less affected by aging, as evident by the high percentage of sugar-ingesting population observed in the aged flies (Figs. 1C-1D, S1D-S1E). Presumably, the feeding decision to consume the sweet sugar over water appears to be simpler than the decision relied on the capability to differentiate nutritional values of sugar solutions. We like to consider these two feeding decisions as simple and complex VBFD, respectively. Taken together, our results suggest the age-dependent deterioration of VBFD may not be directly linked to the loss of sweet sensation. Instead, the impaired VBFD may be associated with the decline of cognitive function.

**Fig. 1.**
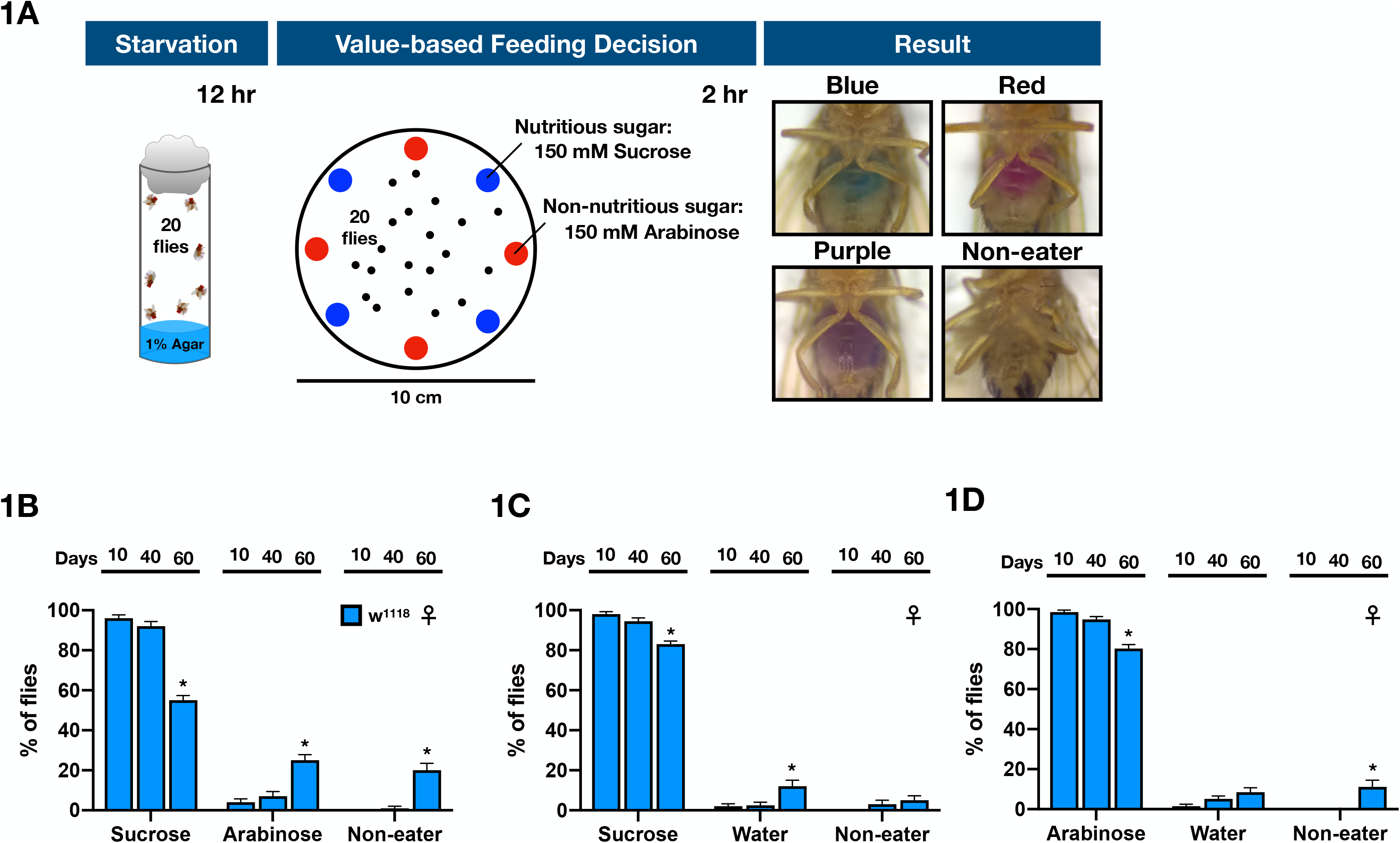
The efficacy of making proper VBFD is gradually deteriorating in aging flies. **(A)** A schematic illustration of VBFD assay. Briefly, food-deprived flies (12 hours of starvation) were presented with two liquid food choices and allowed to feed for 2 hours at 25°C. Each liquid food is labeled by a red or a blue dye. The feeding decision was scored by examining the colors shown in the abdomen. Red or Blue color shown in the abdomen suggests flies only ingest only either one food choice. Purple indicates the flies consume both food choices during the feeding assay. Non-eater means the flies have no labeled food in the digestive tract. **(B-D)** VBFD assays were performed in female flies of different chronological ages: 10, 40, and 60 days. The food choices used were 150 mM sucrose and 150 mM arabinose in **(B**); 150 mM sucrose and plain water in **(C)**; 150 mM arabinose and plain water in **(D)**. Results were expressed as means ± SEM and analyzed by two-way ANOVA. n=100 for each condition. *: p<0.01. The statistical significance was assessed by the comparison to 10 day-old flies.

### VBFD in flies that have altered life expectancy

Given the decision making is regarded as a basic cognitive process (*1, 2*), we used the VBFD assay to assess the cognitive status of Drosophila subjected to selected physiological/pathological conditions. For example, there are various genetic manipulations that effectively alter the life expectancy of Drosophila. It is especially interesting to know if the aged long-lived flies could still preserve the functional cognition. Studies by Morrow et al. (*13*) have shown ectopic expression of Hsp22 (heat shock protein 22) significantly increases both the median and maximal lifespan of fruit flies. Here, the Hsp22-mediated long-lived flies were assayed to determine their efficacy of VBFD making. Our survival analysis indicated the median lifespan of Hsp22-expressing female flies was ~108 days (Fig. 2A). Therefore, VBFD assays were performed in 10 day and 70 day-old female flies. As the population of sucrose-ingesting flies declined drastically in the 70 day-old control flies, ~80% of the age-matched long-lived flies still made the proper VBFD by ingesting only sucrose, but not the arabinose (Fig. 2B). However, the efficacy of proper complex VBFD was eventually declined in the 90 day-old long-lived female flies (Fig. S2A). Less than 60% of 90 day-old female long-lived flies chose the sucrose, albeit this percentage was still higher than 70 day-old control flies. Therefore, our results suggest the pro-longevity effects elicited by ectopic Hsp22 partly slow down the aging-related cognitive decline. In comparison to the VBFD between nutritious and non-nutritious sugars, feeding decisions between the sweet sugar and plain water, which could be considered as a simpler form of VBFD, were less affected in both the age-matched control and long-lived female flies (Figs. 2C-D; results of male flies were shown in Figs. S2D-S2G). Nonetheless, the fact that 90 day-old long-lived female flies were unable to differentiate the sweet sugars from the plain water suggests the sweet sensation may not be fully functional even in the presence of pro-longevity effects (Figs. S2B-S2C). Together, our results indicate the Hsp22-elicited pro-longevity effects are able to improve the maintenance of cognition in aged flies. Furthermore, contrary to the pro-longevity effects, down-regulating Hsp22 expression through the developmental stages drastically reduced the life expectancy of flies (Fig. 2A; UAS-miR-hsp22). Intriguingly, while the complex VBFD was jeopardized in the 10 day-old Hsp22-knock down female flies, the recognition of sweet sucrose and arabinose remained intact, suggesting this anti-longevity effects have a significant impact on the complex VBFD but not the basic sweet sensation (Figs. 2B-D). In addition to the genetic manipulations, lithium chloride (LiCl)-treated long-lived flies were also able to maintain the proper VBFD when they are aged (*14*) (Fig. 2E). In summary, the two pro-longevity treatments we tested not only extend the life expectancy, but also have the capability to preserve the cognitive function in aged flies.

**Fig. 2.**
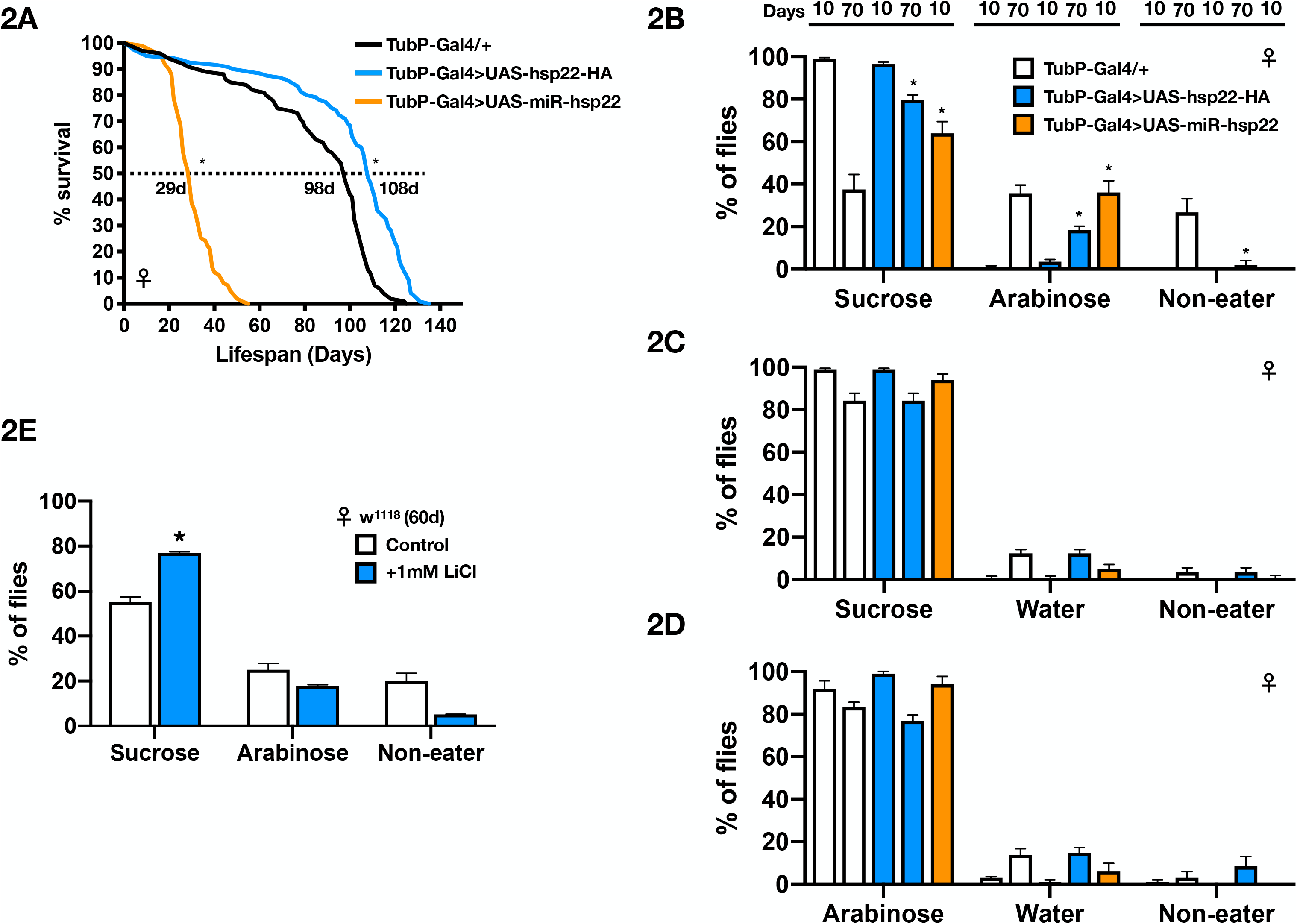
Aged long-lived flies still maintain the proper VBFD. **(A)** Survival curves of Hsp22 over-expressing (OE; UAS-hsp22-HA) and knock-down (KD; UAS-miR-hsp22) flies. Results were analyzed by the log-rank test. n=100 for each genotype. *: p<0.01. **(B-D)** VBFD assays were performed in female flies with altered life expectancy mediated by up- or down- regulating the expression of Drosophila Hsp22. Flies of two chronological ages were used: 10 and 70 days. The food choices used were 150 mM sucrose and 150 mM arabinose in **(B)**; 150 mM sucrose and plain water in **(C)**; 150 mM arabinose and plain water in **(D)**. Results were expressed as means ± SEM and analyzed by two-way ANOVA. n=100 for each condition. *: p<0.01. The statistical significance was assessed by the comparison to age-matched controls. **(E)** Female w^1118^ flies fed with 1mM LiCl for 60 days were given the choices between 150 mM sucrose and 150 mM arabinose in the VBFD assay. Results were expressed as means ± SEM and analyzed by two-way ANOVA. The statistical significance was assessed by the comparison to age-matched controls. *: p<0.01. n=100 for w^1118^ w/o LiCl and n=40 for w^1118^ w/ LiCl. Note that the column of w^1118^ w/o LiCl treatment was the same as presented in Fig. 1B. Genotypes: control (TubP-Gal4/+); OE (UAS-hsp22-HA/+; TubP-Gal4/+); KD (UAS-miR-hsp22/+; TubP-Gal4/+).

### VBFD in flies that have neurodegenerative disorders

Next we like to know whether VBFD is affected in the fly model of neurodegenerative disorders. In this study, the polyglutamine-(polyQ-)induced toxicity was used to promote neurodegeneration (15). The 41Q-HA polypeptides, which contain the HA-tagged 41 poly-glutamine residues, were expressed specifically in adult neurons using the TARGET expression system (*16*). Survival analysis indicated the median lifespan of 41Q-expressing female flies was ~23 days, almost half of the control flies in 29°C (Fig. 3A). Accordingly, VBFD assays were performed in 5, 10, and 15 day-old diseased female flies and the 41Q expression was verified by anti-HA staining (Fig. 3B). As expected, adult flies expressing 41Q for 5 and 10 days still exhibited proper complex VBFD as the polyQ aggregates were gradually accumulating (Fig. 3C). However, after the expression of 41Q for 15 days, only ~50% of the starved diseased flies chose the calorie-rich sucrose. The rest either only ingested the non-metabolizable arabinose or did not consume any food. For the efficacies of simple VBFD, as the expression of 41Q increased, more and more hungry diseased flies chose the non-nutritious water, suggesting the deterioration of simple VBFD and likely the impairment of sweet sensation (Figs. 3D-3E). Taken together, these results suggest the efficacy of proper VBFD is progressively deteriorating in polyQ flies. More importantly, additional to the classical associative learning and memory assays, which involve the pre-conditioning steps, the VBFD analysis could be adapted as an easy and robust assay to determine and quantitate the cognitive disability in diseased flies.

**Fig. 3.**
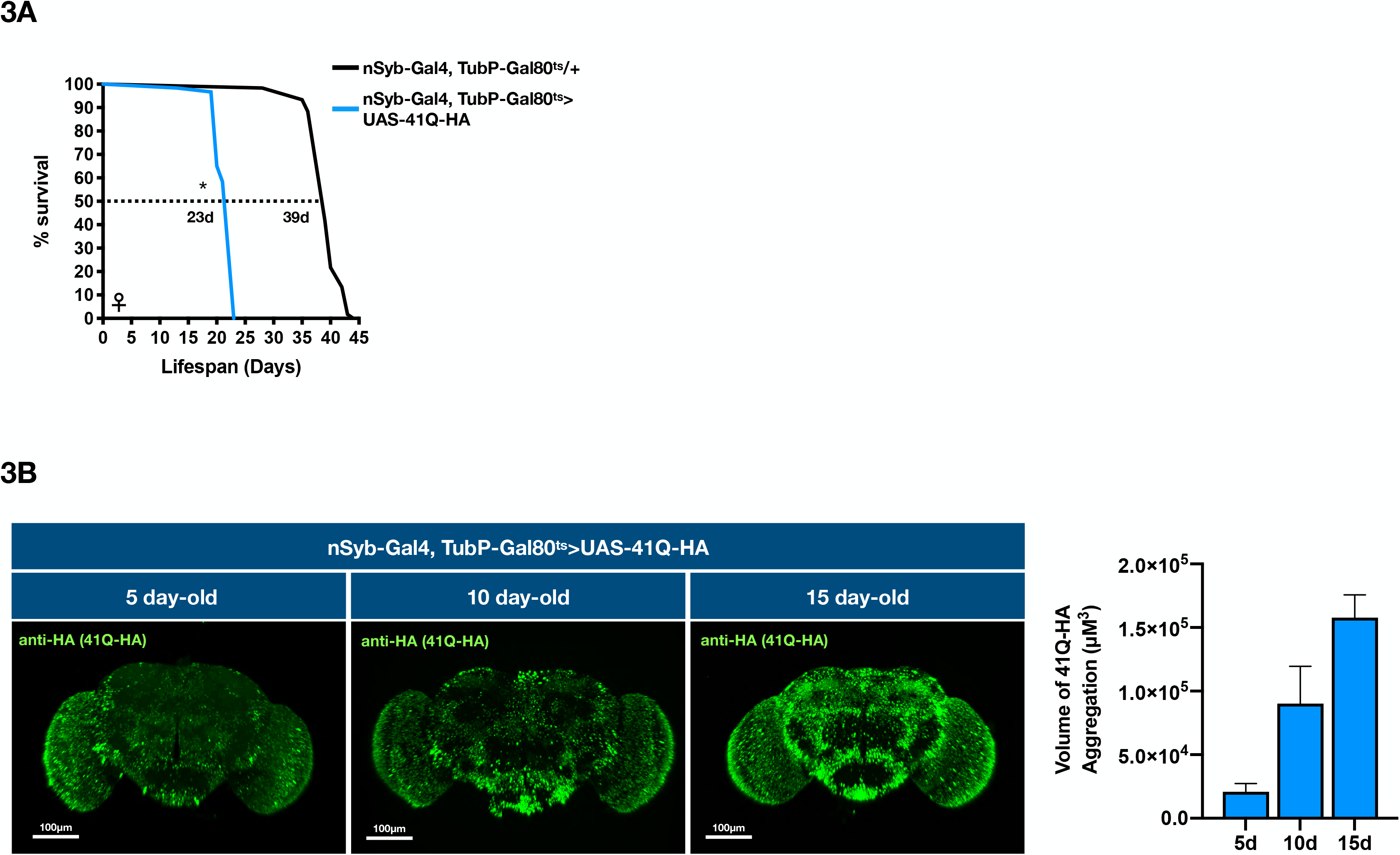
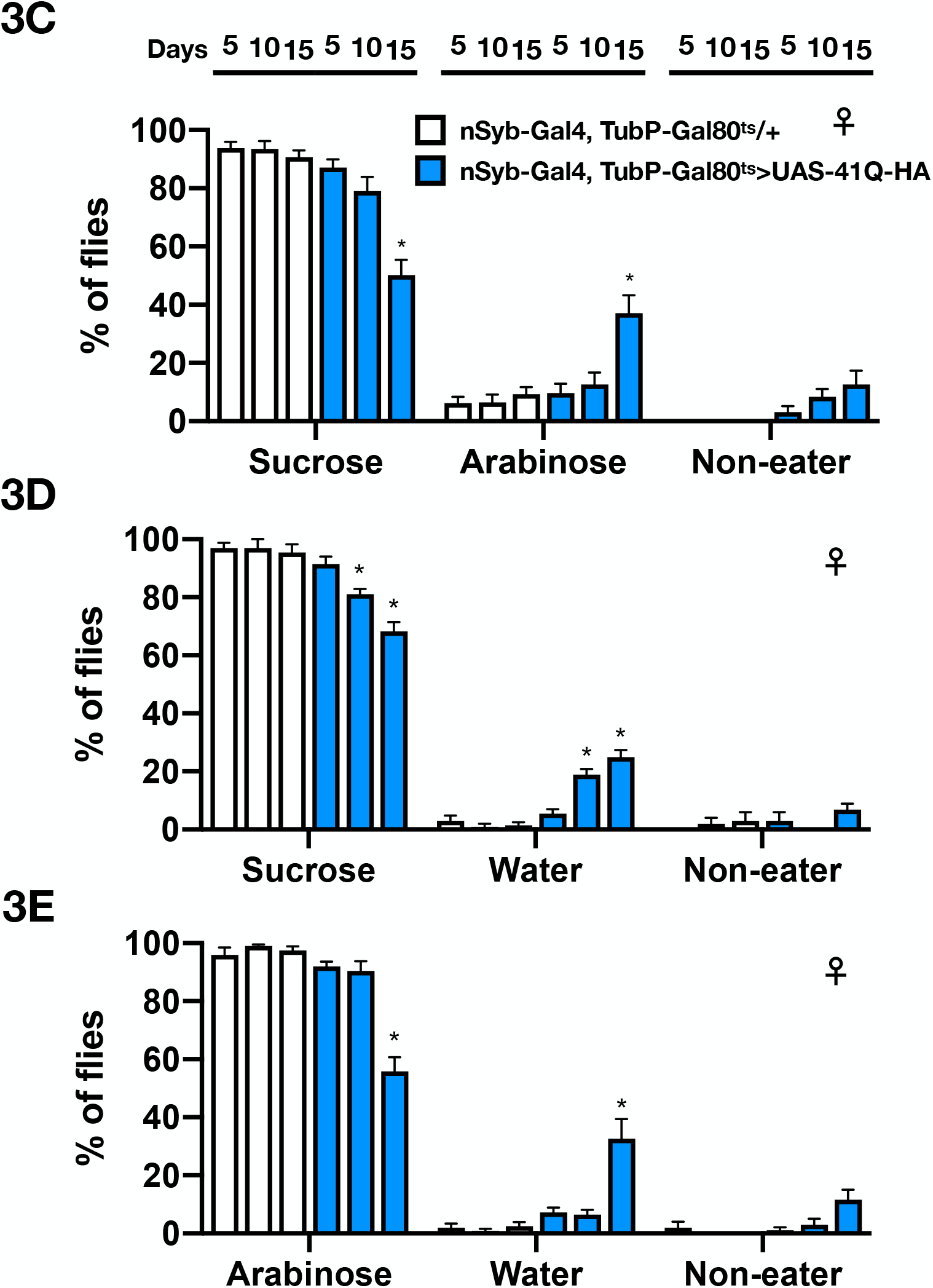
The efficacy of making proper VBFD is impaired in the polyQ-expressing flies. **(A)** Survival curves of flies with adult-onset ectopic expression of 41Q-HA in the nervous system. Results were analyzed by the log-rank test. n=60 for each condition. *: p<0.01. **(B)** Brain images of female flies expressing 41Q-HA for 5, 10, and 15 days. Brain samples were stained with anti-HA (green, stained for 41Q-HA). Scale bars, 100 μm. The volume of 41Q-HA aggregates was quantified as described in the Materials and Methods. **(C-E)** VBFD assays were performed in female flies of different chronological ages (5, 10, and 15 days). The food choices used were 150 mM sucrose and 150 mM arabinose in **(C)**; 150 mM sucrose and plain water in **(D)**; 150 mM arabinose and plain water in **(E)**. Results were expressed as means ± SEM and analyzed by two-way ANOVA. The statistical significance was assessed by the comparison to age-matched controls. n=100 for each condition. *: p<0.01. Genotypes: control (TubP-Gal80^ts^/+; nSyb-Gal4/+); 41Q-HA-expressing flies (TubP-Gal80^ts^/+; nSyb-Gal4/UAS-41Q-HA).

### Effects of LiCl to the polyQ-mediated impairment of VBFD

A variety of compounds and genetic treatments have been shown to confer protective capability against polyQ-mediated pathologies in the Drosophila model (*17*). The most visible pathological feature is the polyQ aggregates. Chemicals, such as histone deacetylase inhibitors (*18*) and LiCl (*19, 20*), can reduce the polyQ aggregates in affected cells. In addition, these protective compounds also partly mitigate several deleterious phenotypes, including eye degeneration, decreased motor activity, and shortened lifespan (*17*). Despite the therapeutic potential against distinct polyQ pathologies, the effectiveness of drug administration to alleviate the cognitive impairment has not been directly tested, given the lack of convenient assays. Here, the VBFD assay was used to determine the cognitive status of diseased flies treated with LiCl (Fig. 4A). Surprisingly, unlike its protective capability to most polyQ-induced defects examined in the earlier Drosophila studies, the impairment of complex VBFD was not alleviated by the treatment of LiCl (Fig. 4B). Application of either 1 mM or 20 mM of LiCl to 41Q-expressing flies only slightly increased the sucrose-ingesting population, while 50 mM LiCl had no effects. Intriguingly, simple VBFDs were partly rescued by the LiCl treatment, suggesting LiCl moderately attenuates the polyQ-mediated disruption of sweet sensation, but fails to improve the decline of complex VBFD (Figs. S3A-S3B).

**Fig. 4.**
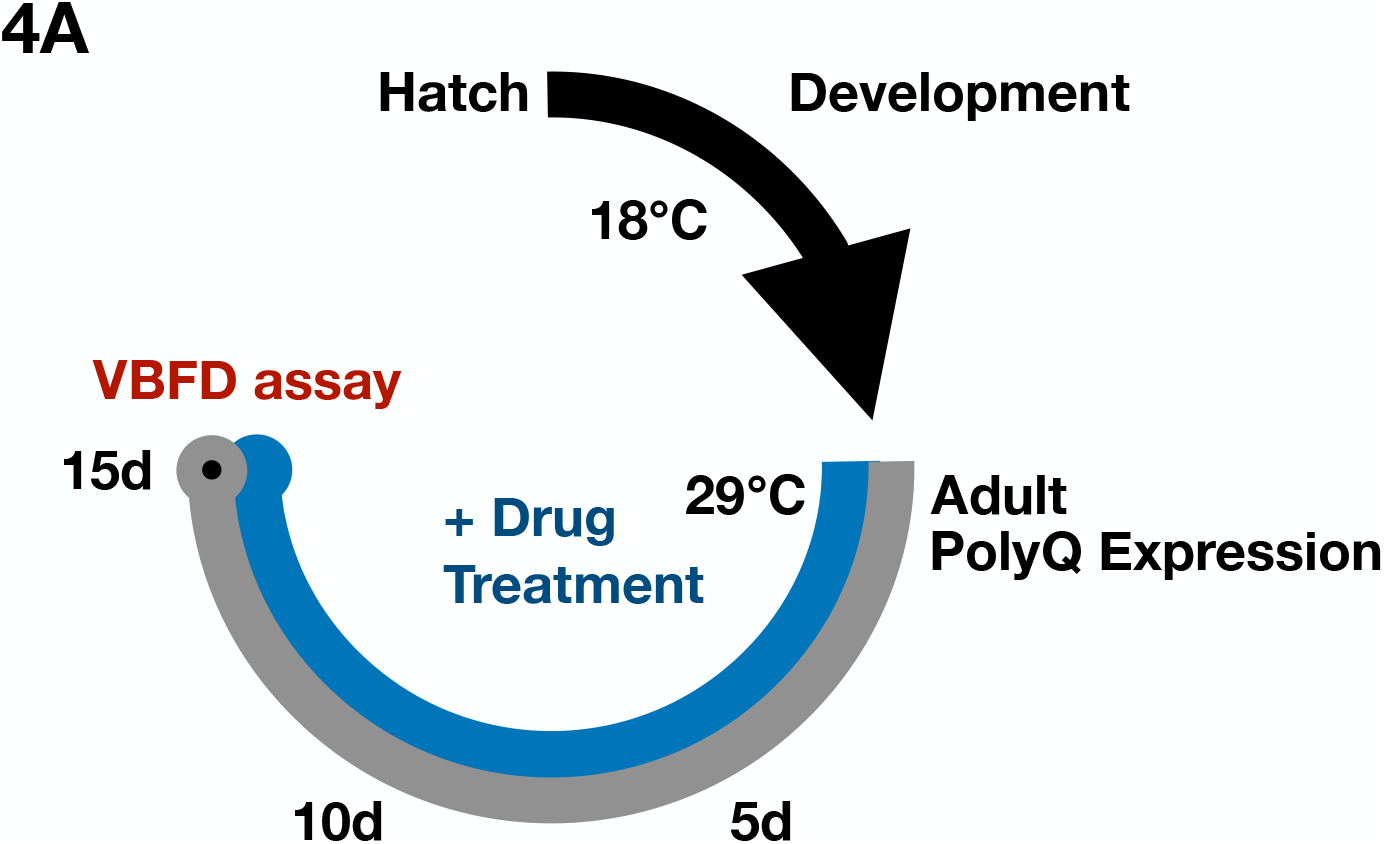
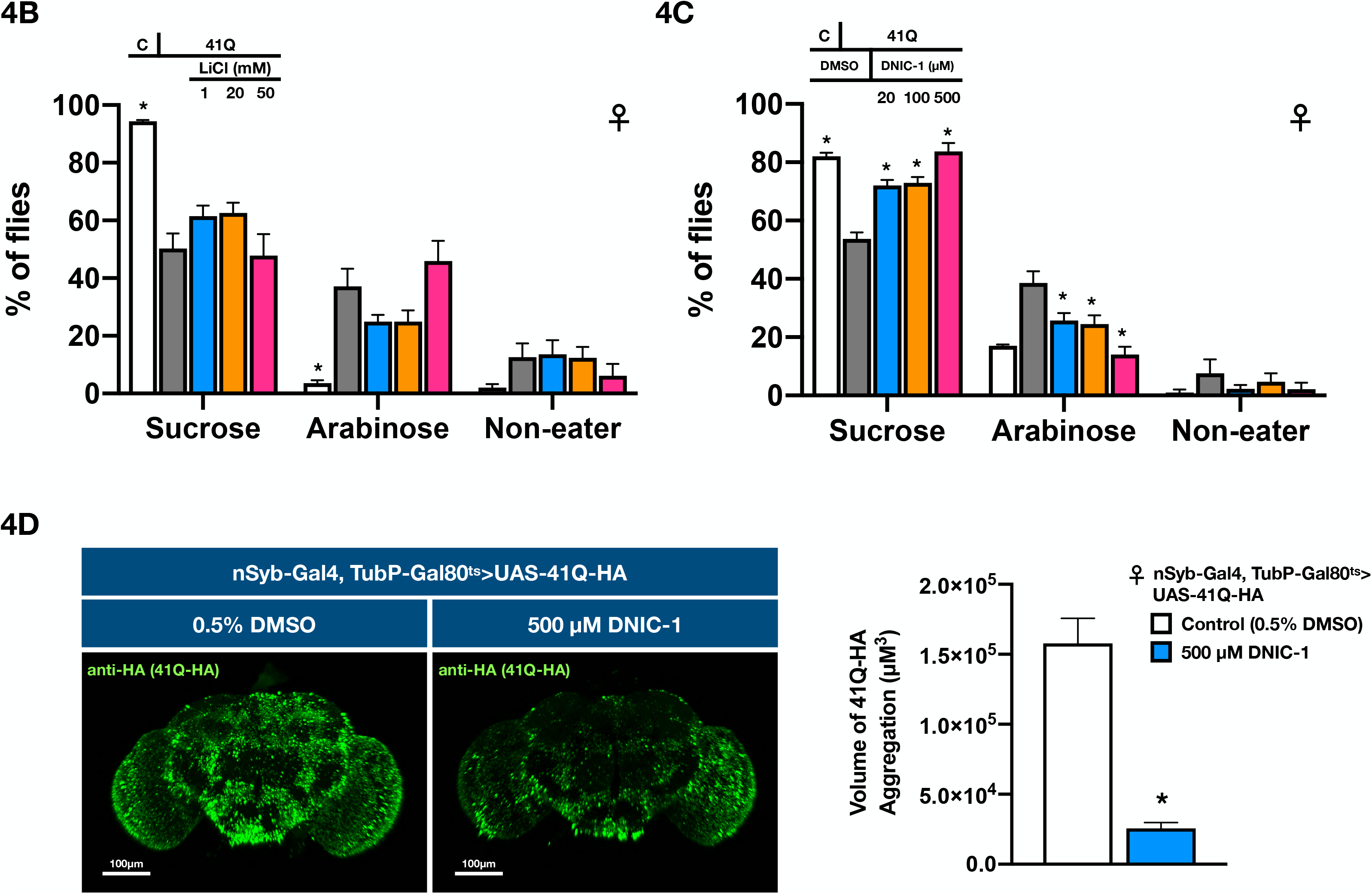
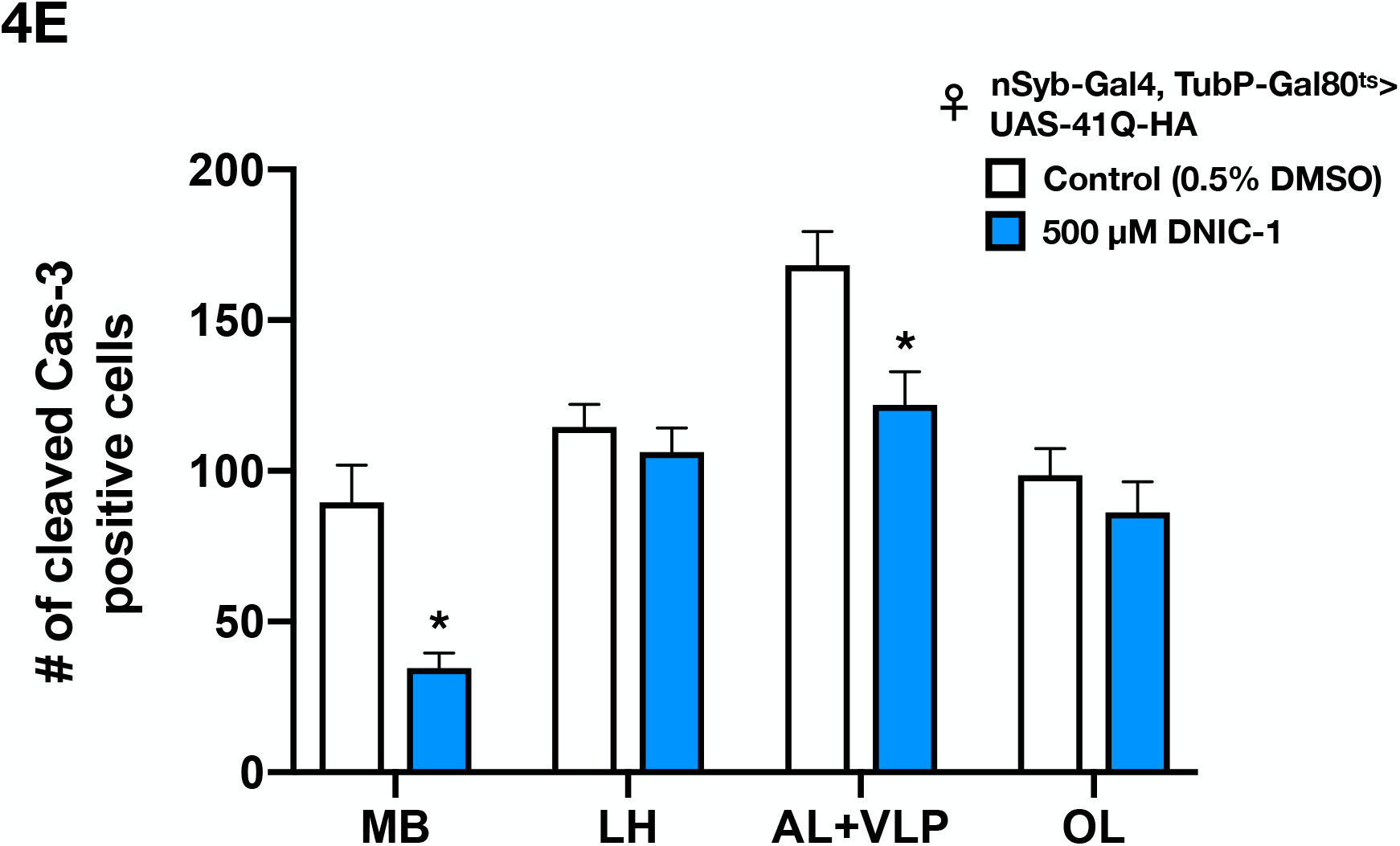
Identification of chemicals that ameliorate the polyQ-induced impairment of VBFD. **(A)** The experimental paradigms of 41Q-HA expression and drug treatments. **(B-C)** VBFD assays (150 mM sucrose vs. 150 mM arabinose) were performed in 15 day-old drug-treated and control female flies. The drugs used were 1, 20, and 50 mM LiCl in **(B)**; 20, 100, and 500 μM DNIC-1 in **(C)**. Results were expressed as means ± SEM and analyzed by two-way ANOVA. n=100 for each condition. The statistical significance was assessed by the comparison to 41Q-HA-expressing flies fed with 0.5% DMSO. *: p<0.01. **(D)** Administration of 500 μM DNIC-1 for 15 days reduced the accumulation of 41Q-HA aggregates in adult brains. Brain samples were stained with anti-HA antibody (green; stained for 41Q-HA). Scale bars, 100 μm. The volume of 41Q-HA aggregates was quantified as described in the Materials and Methods. Results were expressed as means ± SEM and analyzed by Mann-Whitney test. n=10 for each condition. *: p<0.01. **(E)** Administration of 500 μM DNIC-1 for 15 days markedly reduced the number of cleaved Cas-3-positive cells in the mushroom body region of 41Q-HA-expressing female flies. MB, mushroom body; LH, lateral horn; AL+VLP, antenna lobe and ventrolateral protocerebrum; OL, optic lobe. Results were expressed as means ± SEM and analyzed by Mann-Whitney test. n=10 for each condition. *: p<0.01. Genotype for 41Q-HA-expressing flies: TubP-Gal80^ts^/+; nSyb-Gal4/UAS-41Q-HA.

### A novel and steady NO-releasing dinitrosyl iron complex (DNIC-1) ameliorates the polyQ-mediated pathologies

In addition to LiCl, several studies have suggested the neuroprotective properties of nitric oxide (NO) (*21, 22*). To explore the protective functions of NO in the Drosophila model, a novel NO releasing compound, dinitrosyl iron complex [(NO)_2_Fe(μ-SCH_2_CH_2_OH)_2_Fe(NO)_2_] (DNIC-1), was supplemented in the food (*23–26*). As shown in Fig. 4C, DNIC-1 treatment effectively ameliorated the impairment of complex VBFD in 15 day-old female flies suffering from polyQ-mediated neurodegeneration. The fidelity of VBFD could be effectively maintained by the feeding of 500 μM DNIC-1. In addition, the feeding decision-related cognitive function was much less impaired in diseased flies treated with 20 and 100 μM DNIC-1, suggesting a dose-sensitive response (Fig. 4C). As anticipated, the impaired simple VBFD was almost completely preserved by the treatment of 500 μM DNIC-1 (Figs. S4A-4B).

To ascertain the protective effects is NO-dependent, we found the degraded DNIC-1 (use after passing 10x half-lives) was not able to relieve the impairment of VBFD in 41Q flies (Fig. S4G). Moreover, a stable radical scavenger for NO, PTIO, when co-administrated with DNIC-1, markedly abolished the protective function of DNIC-1, substantiating the involvement of NO (Fig. S4G). Besides slowing down the deterioration of VBFD, DNIC-1 treatment also moderately extended the median lifespan of polyQ flies (Fig. S4C; ~18% increase), and restored the eye degeneration and locomotor disability (Fig. S4D and Mov. S1). Together, our results suggest ectopic NO is beneficial to distinct polyQ pathologies. To understand the mechanisms of DNIC-1-mediated neuroprotection, we found levels of polyQ aggregates were greatly reduced by the DNIC-1 treatment (Fig. 4D). The reduction of polyQ aggregates could result from enhanced proteolysis or decreased polyQ synthesis. However, the latter seems less likely given the expression levels of cellular factors, such as Elav and Repo, were not affected by the administration of DNIC-1 (Fig. S4I). It is currently not clear what protein clearance pathways are affected by DNIC-1. Furthermore, in consistent with the reduction of polyQ aggregates, the number of brain cells undergoing cell death was notably decreased in DNIC-1 treated flies (Fig. 4E). Quantification of active Cas-3-positive cells in the selected brain regions indicated the DNIC-1-elicited protective effects were most prominent in the mushroom body (MB), which is a conserved brain structure that plays critical roles in diverse behaviors, including olfaction, associative learning, sleep, and feeding (*27–29*). Therefore, it is possible that loss of MB neurons may interfere with the making of proper VBFD and lead to impaired cognition.

### Neural circuits that govern the efficacy of VBFD

To locate the neural pathways that govern the making of VBFD, we altered the activity of known neuronal types related to the feeding behaviors, starting from the periphery to the central brain. Suppression of neuronal activity was mediated by the expression of Shibire^ts^ (Shi^ts^), a temperature-sensitive mutation of Drosophila dynamin, to block the synaptic transmission in targeted cell types. In consistent with earlier studies, the competence to read the nutritional values and make the proper VBFD was not affected by silencing the activity of gustatory receptor 5a (Gr5a)-positive sweet sensing neurons (*3*) (Figs. 5A, S5A-5B). However, intriguingly, suppression of the Gr66a-positive bitter sensing neurons slightly blunted the complex VBFD (Figs. 5A, S5A-5B). Therefore, as the making of proper VBFD may not require the activity of sweet neurons, the bitter neurons may partly modulate the efficacy of proper VBFD (*3, 30*). Next, in considering the mushroom body (MB) neurons are implicated in diverse aspects of feeding behaviors, such as the integration of hunger/satiety signals and modulation of food-seeking (*29*), the VBFD assay was performed in flies with altered MB activity. Silencing the activity of MB247- or OK107-labeled MB neurons led to moderate changes of VBFD. The sucrose ingesting population was reduced to ~70%, comparing to ~90% of control 10 day-old flies, suggesting the proper VBFD may be partly governed by MB neurons (Figs. 5B, S5C-5D). However, since the VBFD was not fully impaired when MB function was blocked, indicating additional brain regions may participate in VBFD making. However, it is still possible Shi^ts^ proteins may not completely silence the MB activity. Finally, we examined the VBFD in flies with altered dopaminergic rewarding system and feeding-promoting NPF signals (*12, 31, 32*). Surprisingly, the making of proper VBFD remained efficient even when either pathway was perturbed, suggesting other central brain pathways may dictate the proper VBFD (Figs. 5C, S5E-5F). One plausible candidate is the known nutrient sensor in the Drosophila brain, Gr43a (*9*). As shown in Figs. 5D and 5E (more in Figs. S5G-S5J), the efficacy of complex VBFD was not affected by the loss of Gr43a receptors or inhibition of Gr43-positive neurons. Since its functional role in sensing the levels of hemolymph fructose and modulating food consumption, the inability of Gr43a-linked circuits to modulate the efficacy of VBFD further indicates the making of VBFD may depend on multiplexing central brain circuits. Together, our results suggest making proper VBFD may require sophisticated computing processes in the nervous system and MB neurons are a part of the circuits.

**Fig. 5.**
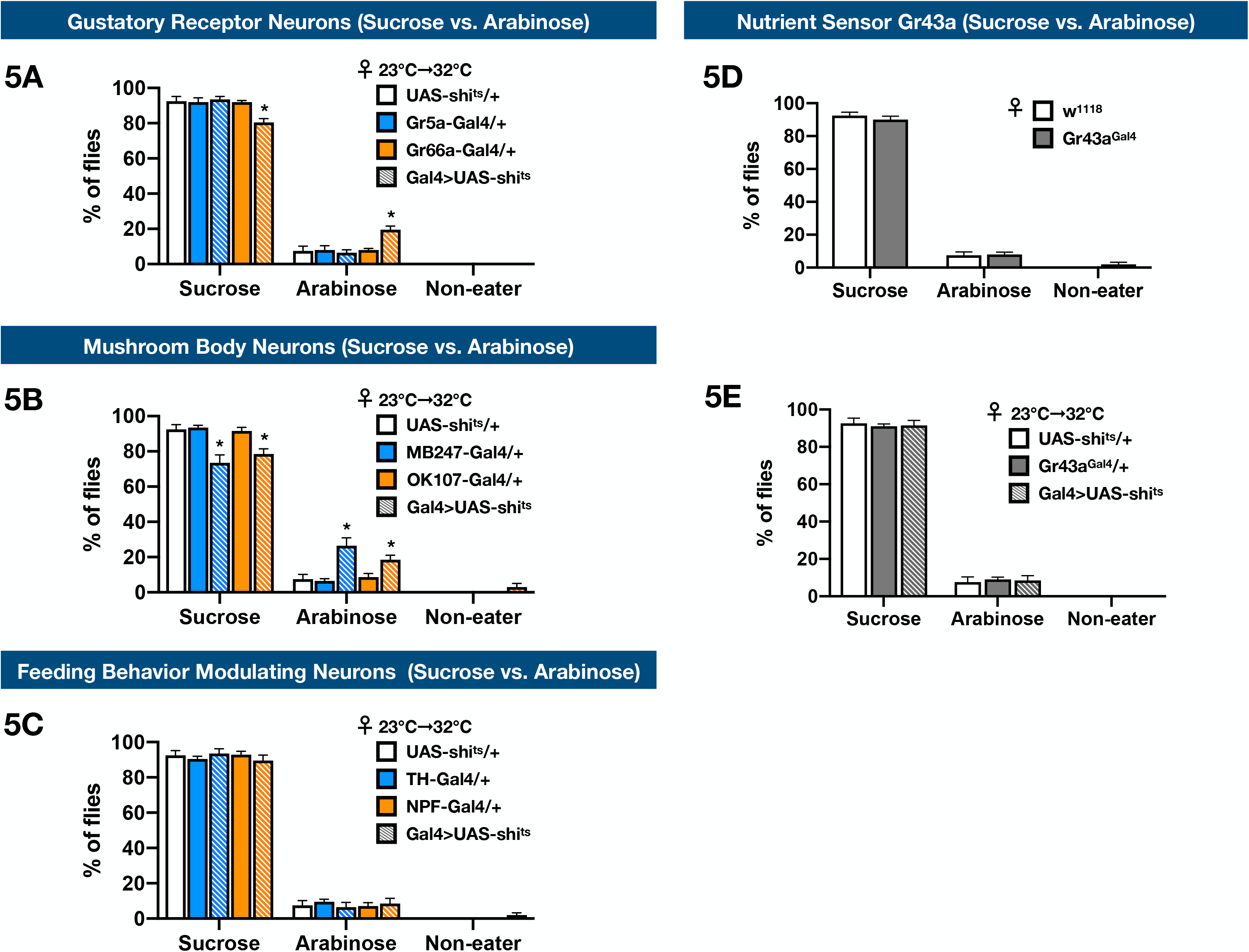
Identification of neural circuits that modulate the making of complex VBFD. **(A-C, E)** VBFD assays (150 mM sucrose vs. 150 mM arabinose) were performed in female flies that have reduced activity in **(A)** gustatory receptor neurons (Gr5a-Gal4 and Gr66a-Gal4), **(B)** mushroom body neurons (MB247-Gal4 and OK107-Gal4), **(C)** feeding behavior modulating neurons (TH-Gal4 and NPF-Gal4), and **(E)** Gr43a-positive neurons (Gr43a^Gal4^). **(D)** VBFD assays were performed in female Gr43a mutants (Gr43a^Gal4^). Results were expressed as means ± SEM and analyzed by two-way ANOVA. n=100 for each condition. Note all the columns of UAS-shi^ts^/+ were the same. In **(A-C, E)**, the statistical significance was assessed by the comparison to UAS-shi^ts^/+ controls. In **(D)**, the statistical significance was assessed by the comparison to w^1118^. *: p<0.01.

## Discussion

Earlier studies have indicated starved fruit flies are able to evaluate and learn the nutritional value of sugar solutions (*3, 6–8*). Particularly, making such VBFD is robust and a native behavior. Consistently we noticed the flies defective of learning and memory, such as Rut and Dnc mutants, could still made proper VBFD (Figs. S5K-S5M). More interestingly, recent studies also indicate the capability of fruit flies to detect nutritional values of sugars involves a taste-independent metabolic sensing pathway (*3, 8*). But how this metabolic sensing pathway finalizes the feeding decision is far from clear. Decision making is an important feature that allows every living animal to identify and choose alternatives based on various criteria, therefore representing a unique spectrum of cognition. Food-deprived flies, once encounter the potential foods, must quickly evaluate the qualities of foods and select the food choice with nutritional value to resolve the energy need. Consequently, the feeding decision based on the caloric contents of foods needs to be properly made and maintained. In this study, we deployed the VBFD assay to determine the cognitive status of Drosophila. Perceivably, the feeding decision between the metabolizable sucrose and non-metabolizable arabinose is more difficult to consolidate than the decision between sweet sugar and plain water, given the differentiation of nutritional values of sugar solutions should be more complicated than the identification of palatable sugar. Along these lines, the VBFD assay and the combinations of testing food substances allowed us to distinguish two distinct levels of cognitive function in the forms of complex and simple VBFDs. By monitoring the efficacy of VBFD, our results showed the deterioration of complex VBFD is more severe in aged flies, while the simple VBFD and perhaps the basic sweet sensation remain less affected (Figs. 1B-1D). Therefore, it appears the decline of complex cognitive functions is more sensitive to the aging process. Moreover, based on the experimental specifications, the complexity levels of feeding decision can be directly characterized.

There are a number of genetic manipulations and chemicals that have pro-longevity effects in the Drosophila model (*13, 14, 33, 34*). However, in addition to the extended life expectancy and improved motor performance in aged flies, their effects to the age-dependent cognition decline has not been systematically explored. In this study, our results showed the efficacies of both complex and simple VBFDs are significantly preserved in the Hsp22-expressing and LiCl-treated long-lived aged flies (Figs. 2B-2E), suggesting, at least in these two cases, the pro-longevity treatments are able to delay the deterioration of feeding-related cognitive behaviors. However, unlike our results, a previous study has shown the efficacy of olfaction-associated learning and memory is not retained in the dietary-restricted long-lived flies (*35*). Since each pro-longevity treatment may exert its effects through distinct mechanisms, it is not clear if other pro-longevity treatments have similar protective effects to the age-dependent cognitive decline. More interestingly, it has been demonstrated in several studies that tissue-specific expression of selected pro-longevity gene is sufficient to prolong the animal’s lifespan (*13, 34, 36, 37*). In that case, learning how these long-lived flies perform in the VBFD assay will allow us to substantiate and characterize the functional inter-connections between the peripheral tissue/organ and the central nervous system, such as the gut-brain communications (*10, 37, 38*).

Cognitive impairment is frequently associated with neurodegenerative diseases. In consistent with this notion, the efficacies of both complex and simple VBFDs were drastically jeopardized in ployQ flies, indicating the development of cognitive disorders and the loss of sweet sensation (Figs. 3C-3E). Unfortunately, the available treatments for neurodegenerative diseases only manage the symptoms or slow down the disease progression. It is also extremely difficult to improve the cognitive impairment associated with neurodegenerative diseases. The neuroprotective effects of LiCl have been demonstrated in several recent studies, including in the SCA3 Drosophila model, which is also a polyQ neurodegenerative disorder (*17, 19, 20*). Chronic administration of LiCl alleviates SCA3-mediated pathologies, such as eye degeneration, locomotor disability, and shortened lifespan. Despite its neuroprotective potential, however, the effect of LiCl to relieve the impairment of cognition has never been examined. Unexpectedly, our VBFD analysis indicated LiCl treatment has only limited effects on the polyQ-mediated impairment of complex VBFD, suggesting the complex cognitive behaviors are more sensitive to the degenerative pathology (Figs. 4B and S3). Regarding the neuroprotective properties of NO, in this study, we found DNIC-1, a novel NO-releasing chemical, is a promising neuroprotective compound that mitigates diverse polyQ-dependent phenotypes, including the decline of cognitive function (Figs. 4C-4E and S4). However, we also noted the protective effects of DNIC-1 is most profound when the compound is administrated along with, but not after, the onset of 41Q expression (Figs. S4E-S4F). In considering the unstable nature and short half-life of NO (~sec-min), selection/development of the prodrug featuring steady and long-term NO-releasing ability is a critical step for the translation/development of NO as a novel therapeutic agent, especially for chronic diseases. At present, there are several classes of NO donor chemical available. In addition to FDA-approved organic nitrates (i.e. glycerol trinitrate), the NONOate variants of NO donor chemical release NO through a hydrolytic mechanism and have distinct half-life (t_1/2_ = 1 min to 20 hrs at pH7.4, 22-25°C).

The design of DNIC-1 [(NO)_2_Fe(μ-SCH_2_CH_2_OH)Fe(NO)_2_], a novel NO-delivery reagent used in this study, is based on a natural motif, dinitrosyl iron unit [Fe(NO)_2_] for the delivery and storage of NO (*23–26*). In comparison to the burst release of NO by the FDA-approved organic nitrates and well-studied NONOates, DNIC-1 displays a steady kinetics for O_2_-triggered release of NO (half-life = 27.4 hrs at pH7.4, 22-25°C), providing a more stable source of ectopic NO under physiological conditions. To verify the neuroprotective effects is mediated by NO, the degraded DNIC-1 was not able to save the impaired VBFD (Fig. S4G). Moreover, feeding of the NO scavenger, PTIO, markedly accelerated the deterioration of VBFD elicited by polyQ aggregates (Fig. S4H). Based on the levels of polyQ aggregates, DNIC-1 treatment was able to effectively reduce the aggregations of polyQ polypeptides and down-regulate the cell death events in the affected brains (Figs. 4D-4E). There are several possible mechanisms, such as activation of abnormal protein clearance machinery, reduction of disease gene expression, and modulation of cellular factors related to cell death/survival (*17, 39, 40*). These possibilities may need to be validated by additional studies. Finally, given its capability to activate the clearance of disease protein aggregates, it will be worth exploring the protective capability of DNIC-1 in other neurodegenerative diseases and in higher animal models.

## Materials and Methods

### Fly Stocks

Flies were reared on regular cornmeal diet and housed in standard conditions. The following laboratory fly lines were used: TubP-Gal4, nSyb-Gal4, OK107-Gal4, MB247-Gal4, TubP-Gal80^ts^; nSyb-Gal4. We obtained the following fly lines from Bloomington Drosophila Stock Center: w^1118^ (BDSC 3605), NPF-Gal4 (BDSC 25681), TH-Gal4 (BDSC 8848), Gr5a-Gal4 (BDSC 57591), Gr66a-Gal4 (BDSC 57670), UAS-shi^ts^ (BDSC 44222), Rut^2080^ (BDSC 9405), Dnc^1^ (BDSC 6020). Gr43a^Gal4^ was obtained from Dr. Hubert Amrein, Texas A&M University (*9*). GMR-Gal4; UAS-41Q-HA and UAS-41Q-HA were obtained from Dr. Horng-Dar Wang, National Tsing Hua University, Taiwan (*41*). In most cases, flies were reared at 25°C. For the Gal4/UAS-shi^ts^ experiments, flies were reared at 23°C until one hour before the VBFD assay (32°C). To express UAS-41Q-HA in adult flies via the TARGET system (*16*), animals were allowed to develop at 18°C and transferred to 29°C right after eclosion.

### Value-based feeding decision assay (based on two-choice feeding assay)

The two-choice feeding assay was performed according to previous studies with minor modifications (*3–5*). Briefly, 20 flies were food-deprived on 1% agar at 12 a.m. for 12 hours. At 12 p.m., flies were mildly anesthetized by CO_2_ and quickly introduced to a 10 cm-petri dish that contains two different liquid substances. Total 8 droplets (4 droplets for each food choice) were evenly placed around the petri dish. Each liquid droplet contains 20 μL solution. The solution was either color labeled with 0.01% erioglaucine disodium salt (blue dye; Acros Organics, Cat# 229730250) or 0.1% Food Red No. 106 (red dye; TCI, Cat# F0143). The dye added was randomly chosen in each round of experiments. In most cases, feeding assays were performed in complete darkness for two hours at 25°C. For the 41Q-HA-expressing flies, feeding assays were conducted at 29°C. For Gal4/shi^ts^ flies, the feeding assays were conducted at 32°C. The feeding decision of fruit flies was observed under the stereomicroscope and scored by color (B: blue [ingesting only the blue food], R: red [ingesting only the red food], P: purple [ingesting both blue and red foods], and N: non-eater [no color accumulation in the abdomen]) accumulated in the abdomen. The results were presented as percentages of (1) (#_B_ + 1/2#_P_)/ #_total_ (number of flies tested), (2) (#_R_ + 1/2#_P_)/#_total_, and (3) #_N_/#_total_. Two sugars were used in the assay: sucrose (Acros Organics, Cat# 177142500) and arabinose (Alfa Aesar, Cat# A10357).

### Generation of UAS-hsp22-HA transgenic line

Total RNA (TRIzol reagent from Sigma, Cat# T9424) from the heads of 20 female w^1118^ flies was used to generate the Drosophila cDNA library (HiScript I First Strand cDNA Synthesis KIT from BIONOVAS, Cat# AM0675-0050). Subsequently, DNA fragments containing the hsp22-HA were PCR amplified from the cDNA library and cloned into the EcoRI and Xhol sites of pUASTattB (GenBank: EF362409.1). Sequences of the DNA insert were verified before transgenesis. The UAS-hsp22-HA was inserted to the attP40 site on the second chromosome for the creation of transgenic line (WellGenetics, Taiwan). Primers used were listed in **Table S1**.

### Generation of UAS-miR-hsp22 transgenic line

The functional stem-loop structure of the artificial mir-based RNAi_Hsp22 miRNA was created through the first primer set_ dme-Hsp22-mir-1 or dme-Hsp22-mir-2 primers by PCR reaction (**Table S1**). This functional stem-loop miRNA was then extended and added flanking sequences with restriction enzyme sites by the second primer set_ Mir6.1_5’EcoRI/BglII and Mir6.1_3’NotI/BamHI primers to get precursor Hsp22 miRNA unit. The BglII and BamHI restriction enzyme sites of precursor Hsp22 miRNA unit were used for assembling of multiple copies to generate the Hsp22-2miRNAs cassette. Finally, the restriction enzyme double digested EcoRI/BamHI-Hsp22-mir1 and BglII/NotI-Hsp22-mir2 were concurrently integrated into the EcoRI and NotI site region of pUASTattB vector (GenBank: EF362409.1) to generate the pUASTattB_miR-Hsp22-2miR plasmid (*42*). The UAS-mir-hsp22 was inserted to the attP40 site on the second chromosome for the creation of transgenic line (WellGenetics, Taiwan). Primers used were listed in **Table S1**.

### Drug treatment

Briefly, for the drug treatment, flies were raised on the regular cornmeal diet containing the selected drug(s). LiCl (Acros Organics, Cat# 199881000) was dissolved in ddH_2_O and diluted to indicated concentrations in the regular fly food. DNIC-1 and PTIO (Sigma, Cat# P5084) were dissolved in DMSO (Acros Organics, Cat# 348441000) and diluted to final concentration(s) in the food. The final concentration of DMSO was kept at 0.5% in all cases. DNIC-1 was synthesized according to previous publications (*23–26, 43*).

### Lifespan assay

Unmated flies were collected and separated by gender. Twenty flies were housed in a vial. In most of our experiments, flies were incubated at 25°C, except for the flies used for adult-specific expression. Flies carrying the TARGET genetic elements were reared at 18°C until eclosion and transferred to 29°C. Every 2-3 days, flies were transferred to a new vial containing fresh medium and number of the dead flies was counted and recorded. Survival analyses were performed using Prism 7 (GraphPad Software). To verify the life-supporting capability of sucrose and arabinose, 1% agar containing 150 mM of selected sugar was used as the sole food source of 1 day-old w^1118^ flies. Flies were transferred to a new vial containing fresh food preparation every 12 hours, and their survival was monitored and recorded.

### Drosophila eye morphology

10 day-old flies of indicated genotypes were anesthetized by CO_2_ and the eye images were captured under the stereo-microscope.

### Immunohistochemistry

Fly brains were dissected and fixed by 4% paraformaldehyde (Electron Microscopy Sciences Cat# 15710) for 25 minutes at room temperature. The brain samples were washed 3 times with PBST (1x PBS and 0.5% Triton X-100) and incubated in PBST overnight at 4°C. Subsequently, the brain samples were incubated with primary antibodies (diluted in PBST plus 5% of normal goat serum) overnight at 4°C. Next, brain samples were washed three times with PBST and incubated with secondary antibodies for another night at 4°C. Finally, the brain samples were mounted using the SlowFade Gold antifade reagent (Invitrogen, Cat# S36936). Brain images were captured by the Leica SP8 confocal microscope system. Imaris (ver 9.1, Bitplane) was used to process and analyze brain images. Primary antibodies used in this study were: rat anti-HA (1:250; Roche 3F10, RRID: AB_2314622), rabbit anti-cleaved Caspase-3 (1:200; Cell Signaling Technology 9661, RRID: AB_2341188), rat anti-Elav (1:100; DSHB, Elav 7E8A10, RRID:AB_528218), and mouse anti-Repo (1:75; DSHB 8D12, RRID:AB_528448). Secondary antibodies used in this study were: goat anti-rat Alexa 488 (1:500; Invitrogen, A11006), goat anti-rabbit Alexa 568 (1:500; Life Technologies, A11036), and goat anti-mouse Alexa 647 (1:500; Life Technologies, A21236).

### Quantification and statistical analysis

Statistical tests were conducted using Prism 7 (GraphPad Software). Results of feeding decision were expressed as means ± SEM and analyzed by two-way ANOVA with multiple comparisons (Dunnett’s test). The survival curves were plotted as Kaplan Meyer plots and the statistical significance was tested using the log-rank test. Quantification of 41Q-HA aggregates in the brain images was performed using the “Surface Function” of Imaris. The brain region used for the quantification was outlined in Fig. S6A. Number of cleaved Caspase-3-positive cells was calculated using the “Spots Function” of Imaris. Fig. S6B outlined the brain regions used to measure the dying brain cells (MB, mushroom body; LH, lateral horn; AL+VLP, antenna lobe and ventrolateral protocerebrum; OL, optic lobe). The quantitative values were expressed as means ± SEM and analyzed by Mann-Whitney test. All the asterisks in statistical tests indicate statistical significance compared to the corresponding controls (*: p<0.01).

## Acknowledgments

We thank Dr. Hubert Amrein for the Gr43a mutant; Dr. Horng-Dar Wang for GMR-Gal4; UAS-41Q-HA and UAS-41Q-HA fly lines. We also thank the Bloomington Drosophila Stock Center (IN, USA) and Fly Core in Taiwan for providing and handling the fly stocks. This manuscript was aided by comments and discussion from Dr. Yan-Hwa Wu Lee and Dr. Tse-Chun Kuo.

## Funding

This work was supported by the Ministry of Science and Technology, Taiwan (MOST104-2628-B-009-001-MY3 to C.-F. K. and MOST107-2320-B-009-003-MY3 to C.-F. K.) and the “Center For Intelligent Drug Systems and Smart Bio-devices (IDS^2^B) “from The Featured Areas Research Center Program within the framework of the Higher Education Sprout Project by the Ministry of Education (MOE) in Taiwan.

## Author contributions

C.-C. Y., T.-W. C, and C.-F. K. designed the research. C.-C. Y. and F.-F. C. performed all the genetic and behavioral assays, as well as the data analysis. Y.-H. H., T.-T. L., and Y.-M. W. formulated and prepared the DNIC-1 chemicals. J.-C. L., P.-L. C., and C.-H. C. generated the UAS-mir-hsp22 construct and characterized the transgenic line. C.-C. Y. and C.-F. K interpreted the results and wrote the manuscript.

## Competing interests

Authors declare no competing interests.

## Data and materials availability

All data are available in the manuscript or the supplementary materials; raw data are available upon request.

## Supplementary Materials

### Supplementary Figure Legends

**Fig. S1.**
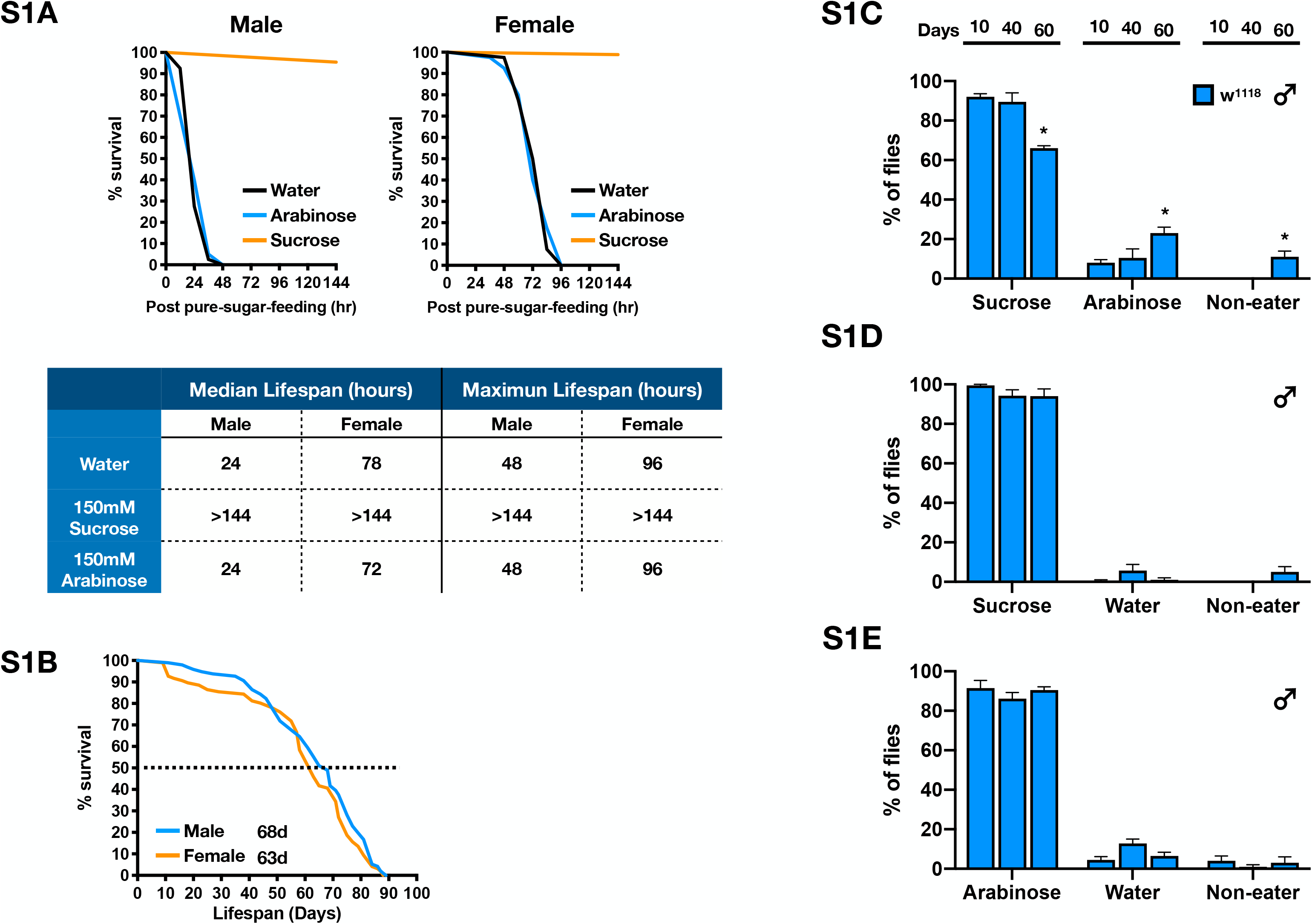
The efficacy of making proper VBFD in the aging flies. **(A)** Survival curves of w^1118^ flies fed on 1% agar (control) and 1% agar plus indicated sugar type (150mM sucrose or 150mM arabinose). n=40 for each condition. **(B)** Survival curves of w^1118^ flies. Median lifespan: female (orange), 63 days, n=97; Male (blue), 68 days, n=96. **(C-E)** VBFD assays were performed in male flies of different chronological ages (10, 40, and 60 days). The food choices used were 150 mM sucrose and 150 mM arabinose in **(C)**; 150 mM sucrose and plain water in **(D)**; 150 mM arabinose and plain water in **(E)**. Results of VBFD were expressed as means ± SEM and analyzed by two-way ANOVA with multiple comparisons (Dunnett’s test). n=100 for each condition. *: p<0.01. Results of 40 day- and 60 day-old flies were compared to 10 day-old flies.

**Fig. S2.**
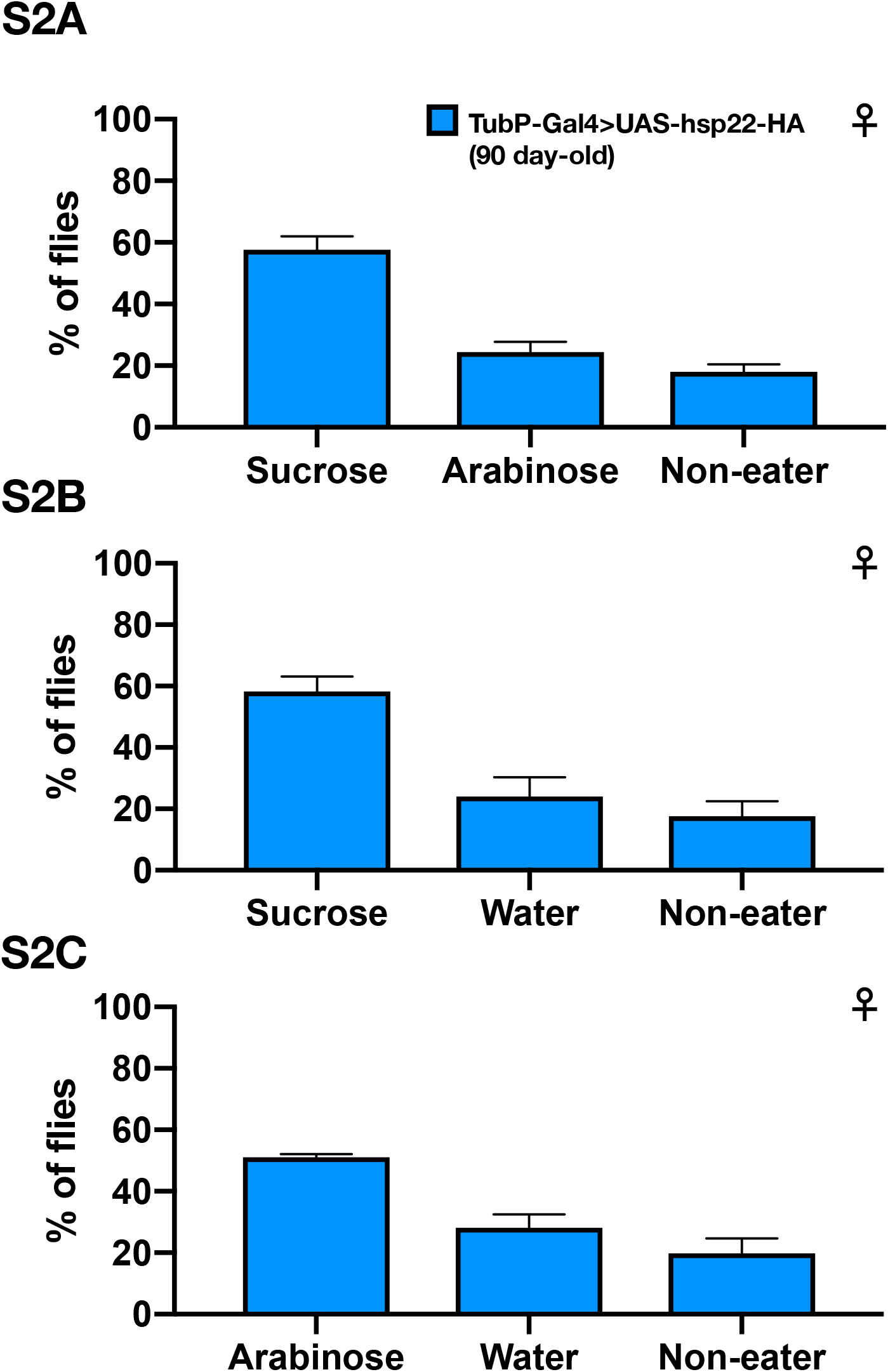
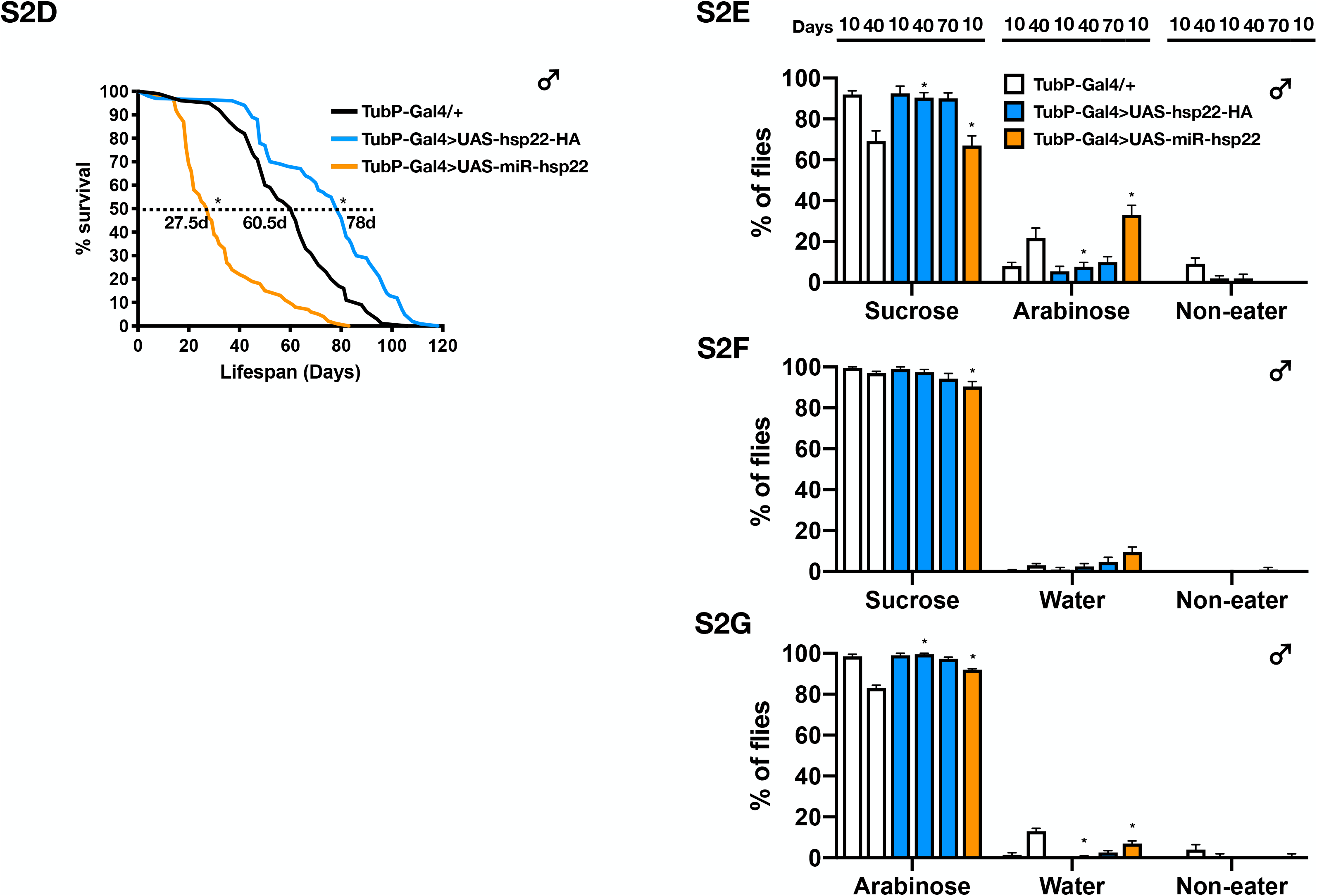
Aged long-lived flies still preserve the proper VBFD. **(A-C)** Results of VBFD assays for 90 day-old female flies expressing ectopic Hsp22. **(D)** Survival curves of Hsp22 over-expressing (OE; UAS-hsp22-HA) and knock-down (KD; UAS-miR-hsp22) male flies. Results were analyzed by the log-rank test. n=100 for each genotype. *: p<0.01. **(E-G)** VBFD assays were performed in Hsp22 OE and KD male flies of different chronological ages (10, 40, and 70 days). The food choices used were 150 mM sucrose and 150 mM arabinose in **(A)** and **(E)**; 150 mM sucrose and plain water in **(B)** and **(F)**; 150 mM arabinose and plain water in **(C)** and **(G)**. Results of all VBFD were expressed as means ± SEM and analyzed by two-way ANOVA with multiple comparisons (Dunnett’s test). n=100 for each condition. *: p<0.01. The statistical significance was assessed by the comparison to age-matched controls. Genotypes: control (TubP-Gal4/+); OE (UAS-hsp22-HA/+; TubP-Gal4/+); KD (UAS-miR-hsp22/+; TubP-Gal4/+).

**Fig. S3.**
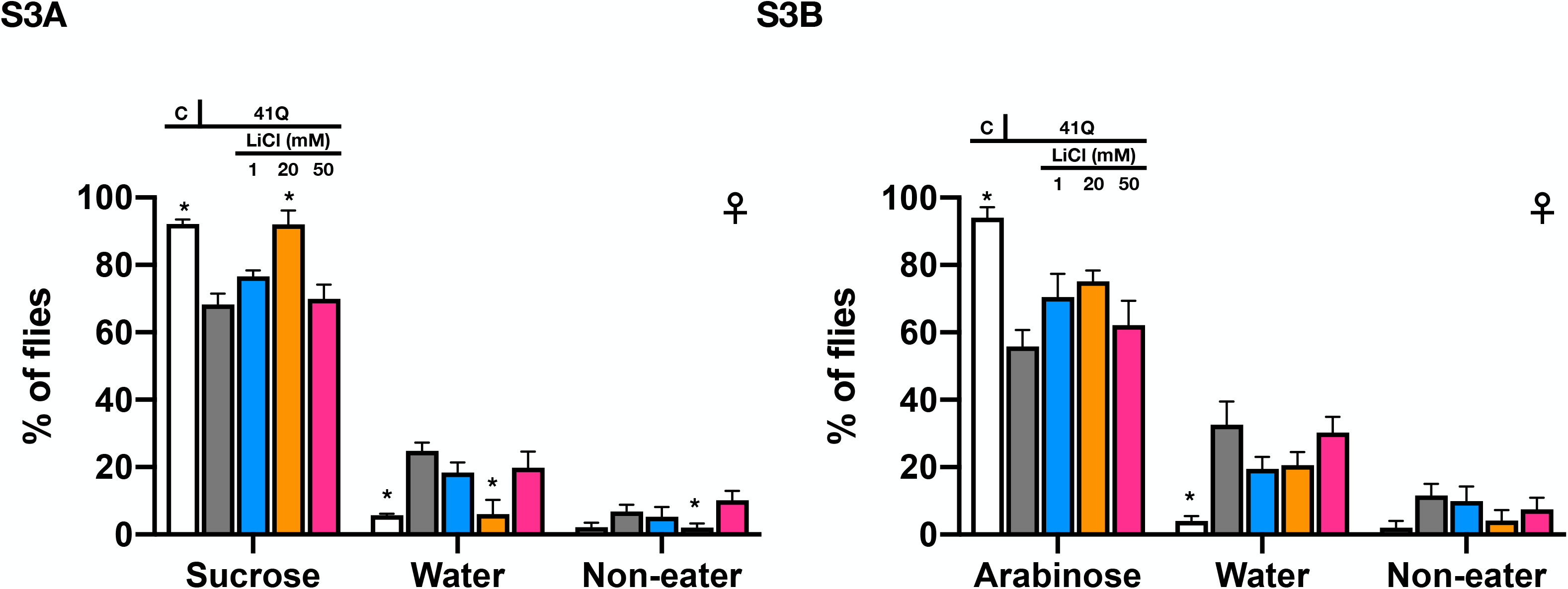
LiCl moderately improves the impairment of simple VBFD caused by polyQ expression. **(A)** and **(B)** VBFD results of 41Q-HA-expressing flies (sucrose vs. water and arabinose vs. water) fed with 1, 20, and 50 mM LiCl. Results were expressed as means ± SEM and analyzed by two-way ANOVA with multiple comparisons (Dunnett’s test). The significance of differences was compared to 41Q-HA-expressing flies without LiCl treatment (gray-colored bars). n=100 for each condition. *: p<0.01. Genotypes: control (TubP-Gal80^ts^/+; nSyb-Gal4/+); 41Q-HA-expressing flies (TubP-Gal80^ts^/+; nSyb-Gal4/UAS-41Q-HA).

**Fig. S4.**
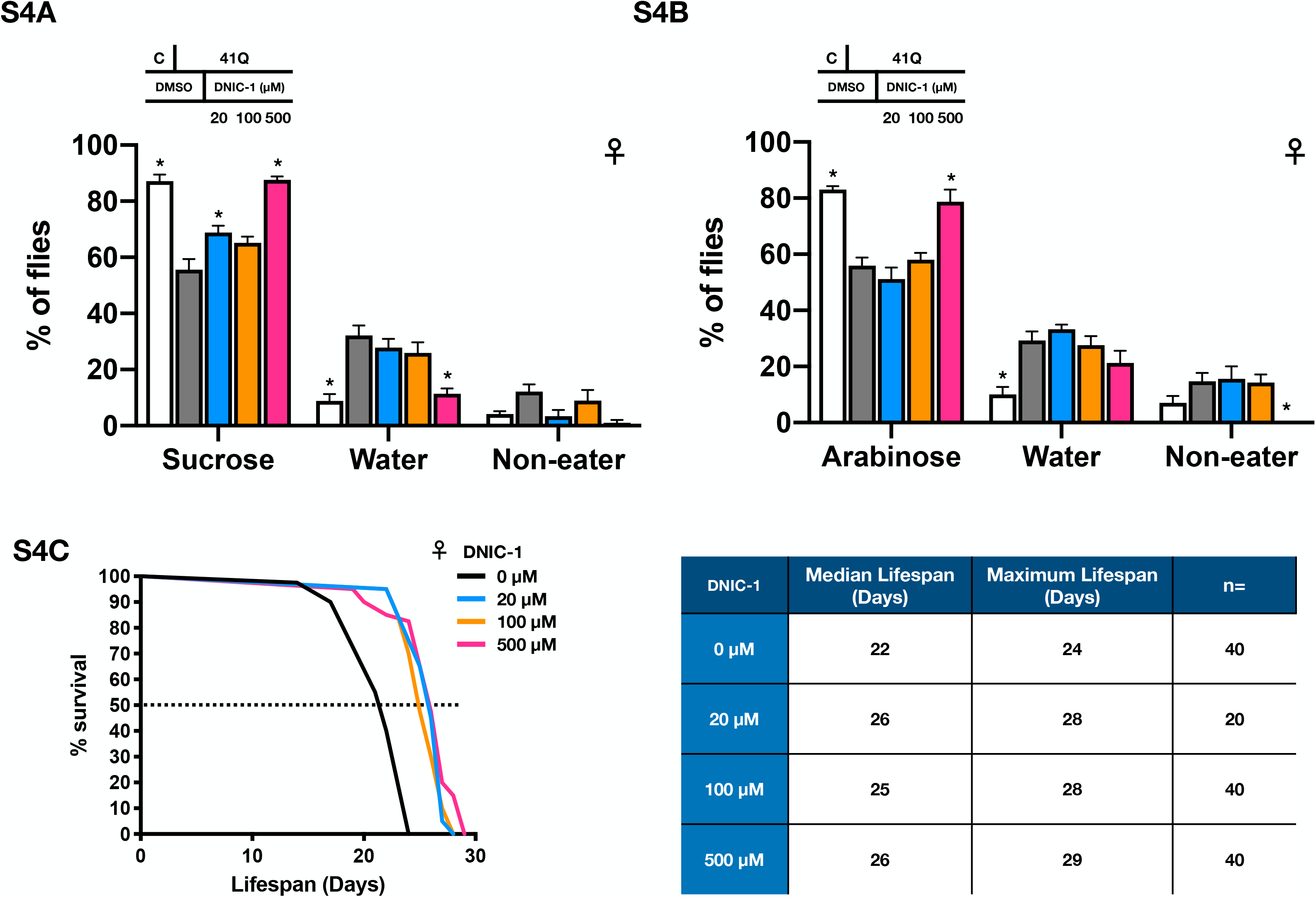
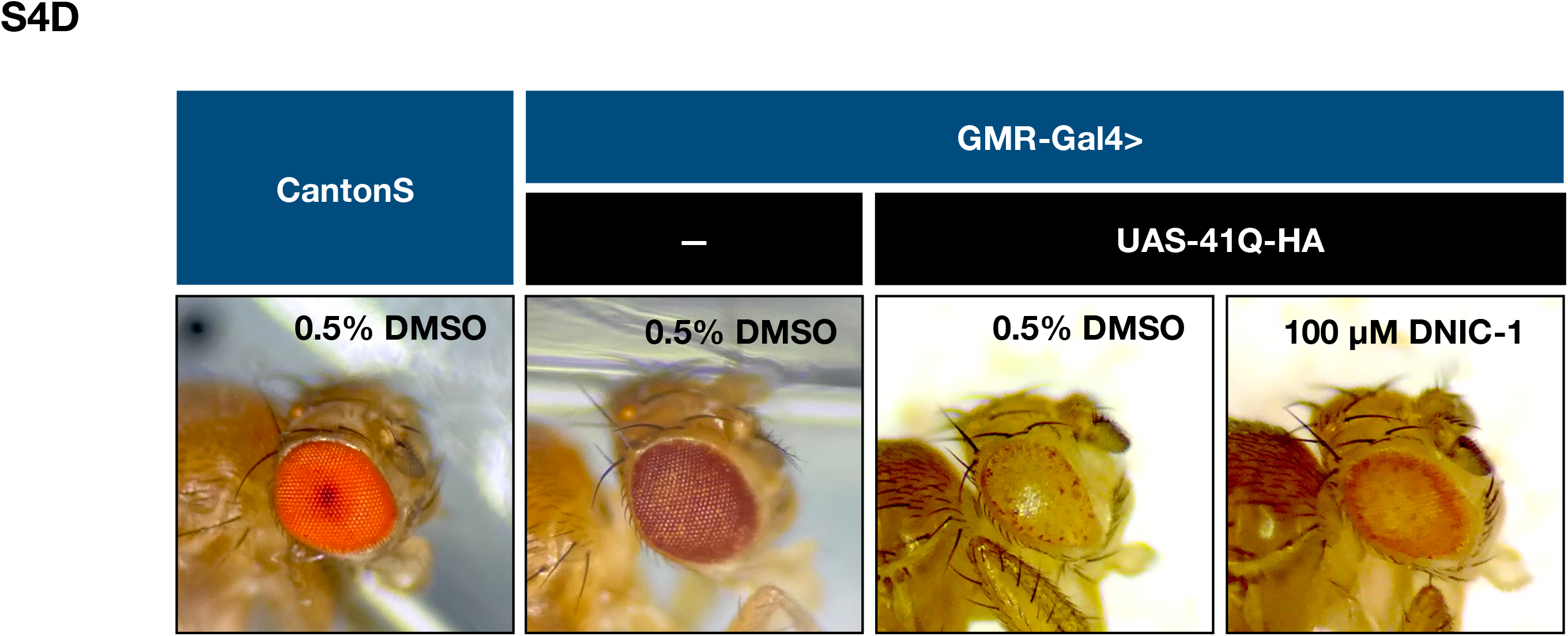
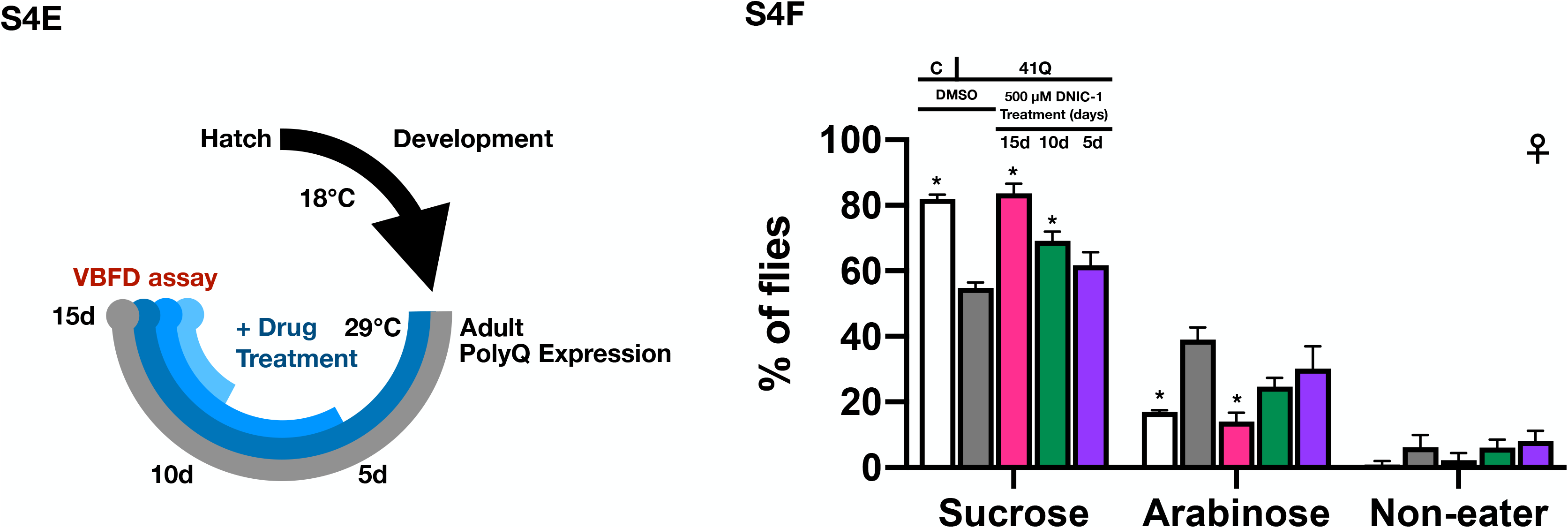
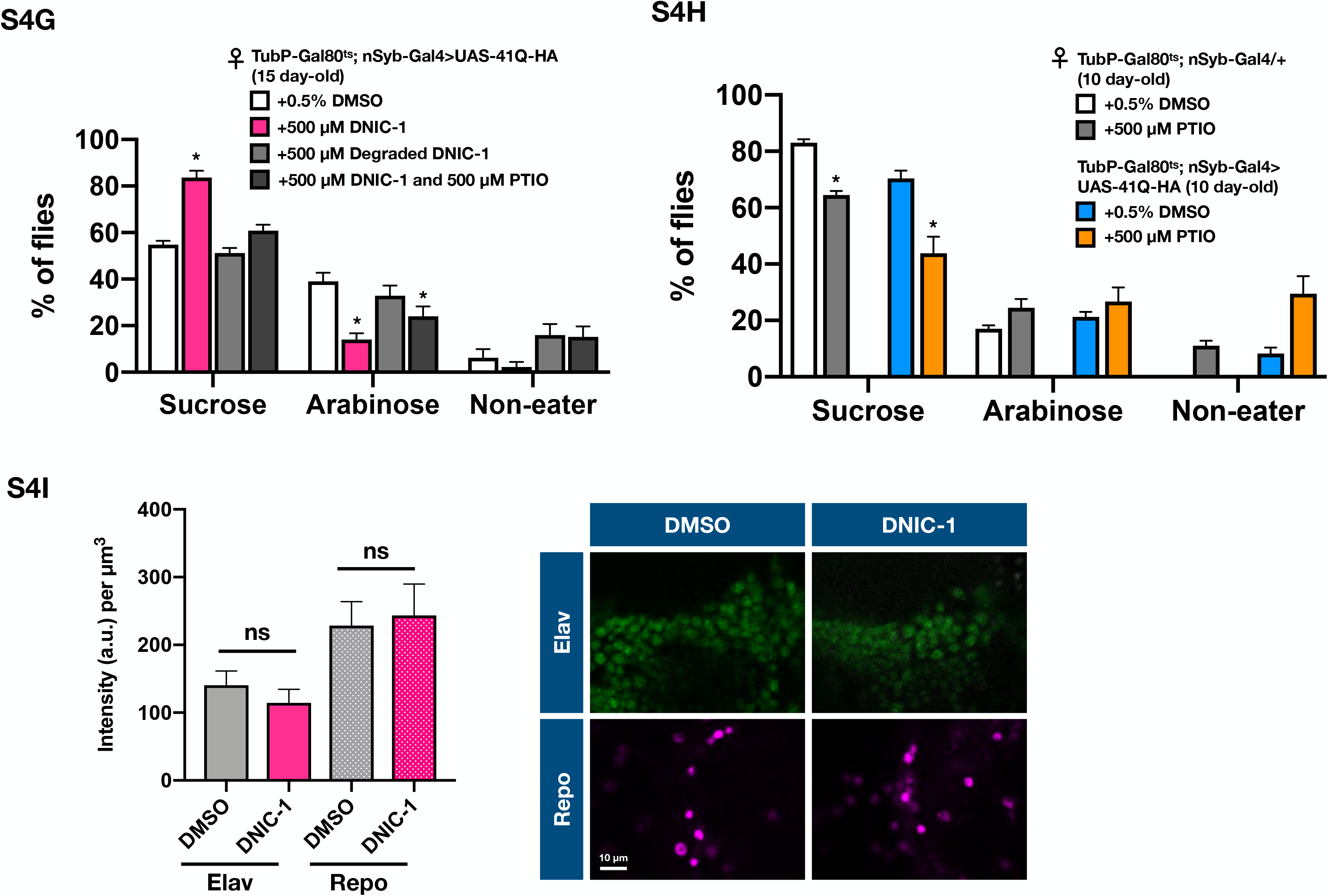
DNIC-1 treatment provides protection to the polyQ-mediated pathologies. **(A)** and **(B)** VBFD results of 41Q-HA-expressing flies (sucrose vs. water and arabinose vs. water) fed with 20, 100 and 500 μM DNIC-1. **(C)** Survival curves of 41Q-HA-expressing female flies fed with 20, 100, and 500 μM DNIC-1. **(D)** Eye images of 41Q-HA-expressing flies fed with 0.5% DMSO and 100 μM DNIC-1. **(E)** The experimental paradigms of 41Q-HA expression and DNIC-1 treatment. In this paradigm, three time points of drug treatment were indicated (total treatment days are 5, 10, 15 days, respectively). Please also note the duration (15 days) of 41Q-HA expression were kept the same for all conditions. Results of VBFD assays after the DNIC-1 treatment were shown in **(F)**. **(G)** VBFD assays were performed in 41Q-HA-expressing flies fed with 500 μM decayed DNIC-1 (after passing 10x half-lives) and 500 μM DNIC-1 + 500 μM PTIO (NO scavenger) for 15 days. **(H)** Results of VBFD assays on 41Q-HA-expressing flies fed with 500 μM PTIO for 10 days. Note that the columns of control flies fed with 0.5% DMSO and 41Q-HA-expressing flies fed with 0.5% DMSO and 500 μM DNIC-1 were the same as presented in Fig. 4C. Results of all VBFD assays shown above were expressed as means ± SEM and analyzed by two-way ANOVA with multiple comparisons (Dunnett’s test). n=100 for each condition. The statistical significance was assessed by the comparison to respective controls. *: p<0.01. Genotypes: control (TubP-Gal80^ts^/+; nSyb-Gal4/+); 41Q-HA-expressing flies (TubP-Gal80^ts^/+; nSyb-Gal4/UAS-41Q-HA). **(I)** Immunostaining signals of Elav and Repo in the brain samples derived from control flies treated with 0.5% DMSO or 500 μM DNIC-1. n=5 for each condition. Quantifications of signal intensity were expressed as means ± SEM, and analyzed by Mann-Whitney test. Genotype: TubP-Gal80^ts^/+; nSyb-Gal4/+. Scale bar: 10 μM.

**Fig. S5.**
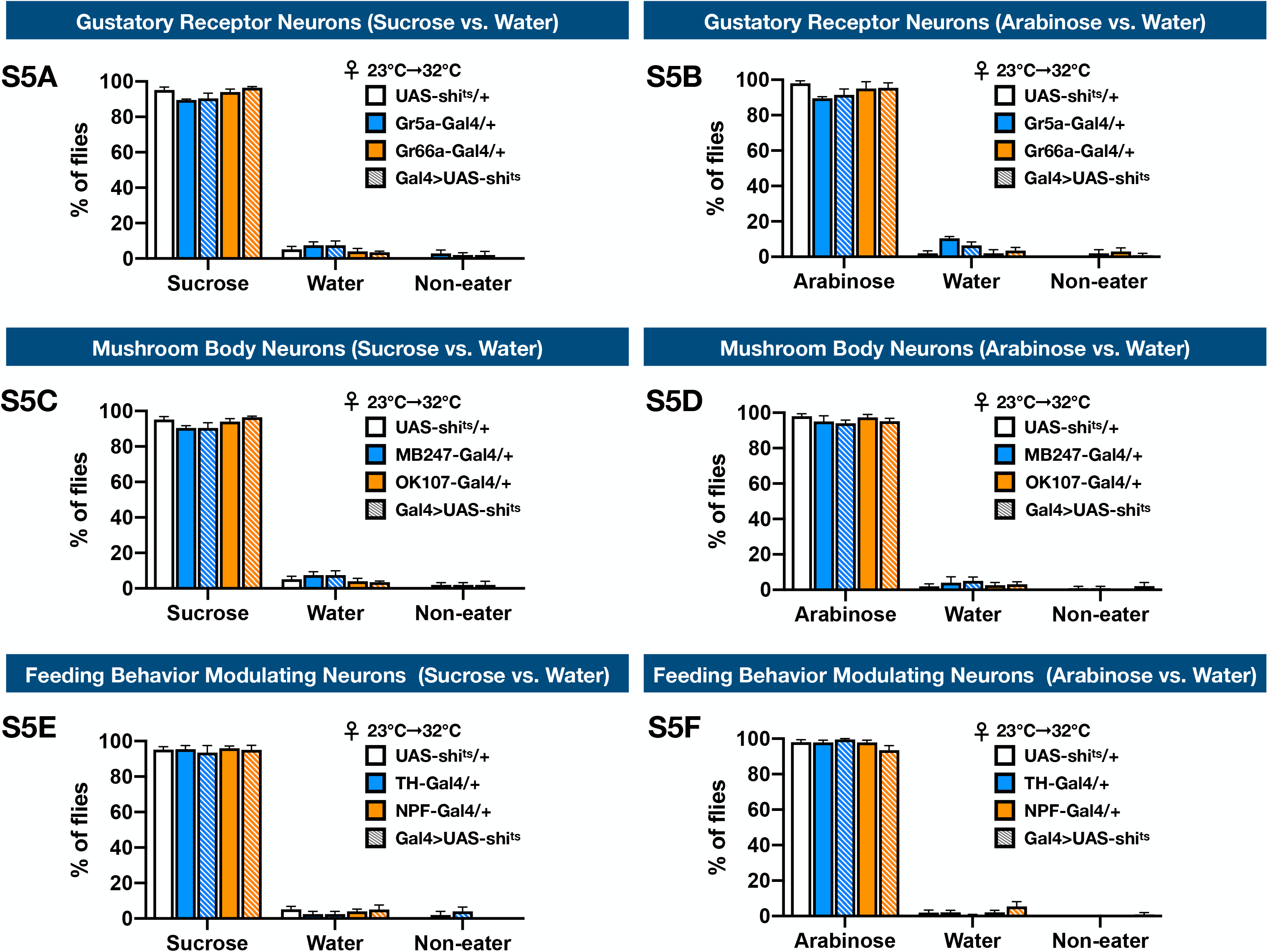
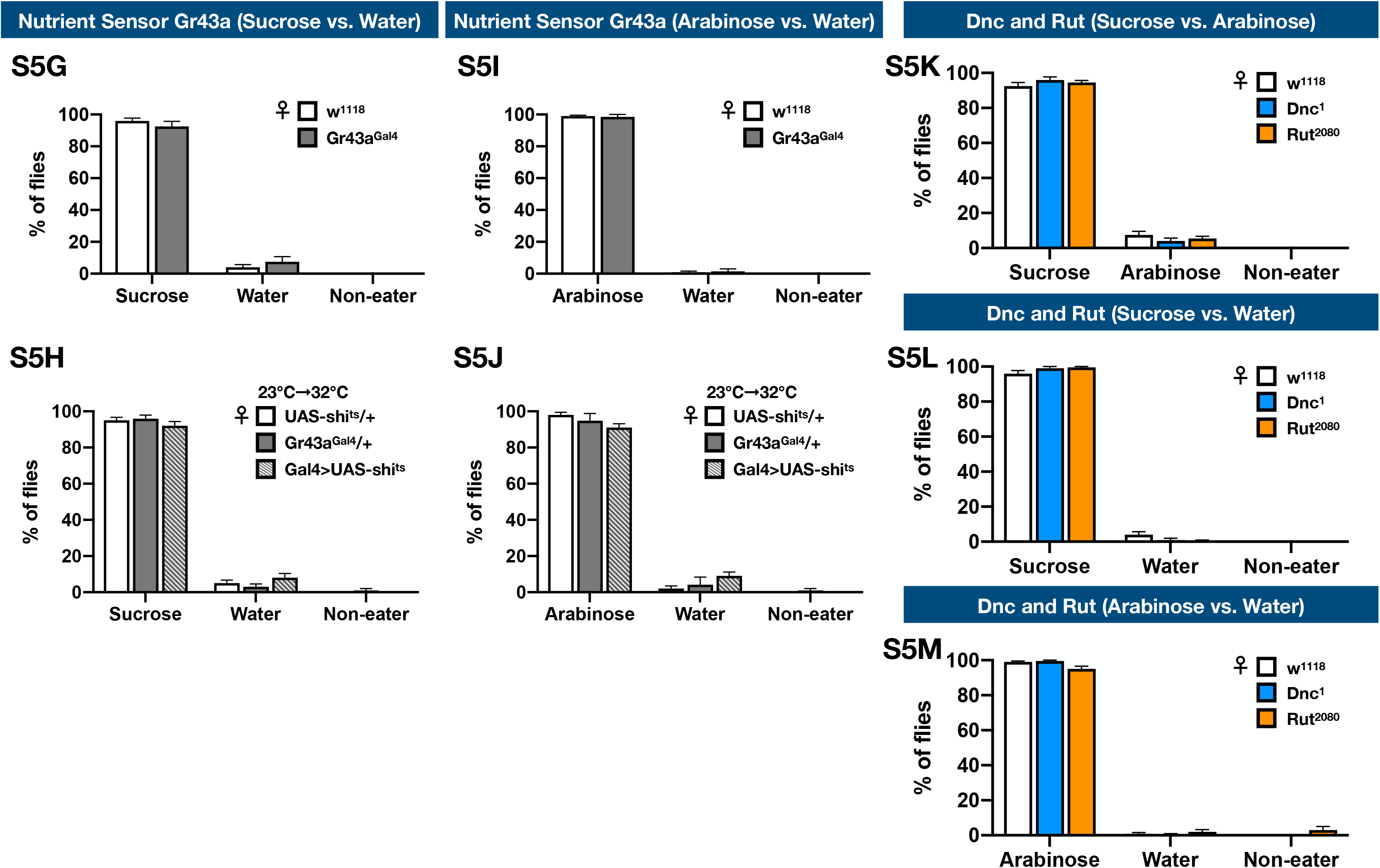
Activity of mushroom body is involved in making the complex VBFD. **(A-F, H-J)** VBFD assays (sucrose vs. water and arabinose vs. water) were performed in female flies that have reduced activity in **(A)** and **(B)** gustatory receptor neurons (Gr5a-Gal4 and Gr66a-Gal4); **(C)** and **(D)** mushroom body neurons (MB247-Gal4 and OK107-Gal4); **(E)** and **(F)** feeding behavior modulating neurons (TH-Gal4 and NPF-Gal4); and **(H)** and **(J)** Gr43a-positive neurons (Gr43a-Gal4). **(G-I, K-M)** VBFD assays were performed in female Gr43a^Gal4^ mutants **(G-I)**, Dnc, and Rut **(K-M)** mutants. 7~10-day old Gal4>UAS-shi^ts^ flies were used. 10 day-old Gr43a^Gal4^, Dnc, and Rut female mutants were assayed. Results of all VBFD were expressed as means ± SEM and analyzed by two-way ANOVA with multiple comparisons (Dunnett’s test). n=100 for each condition. Note that the columns of w^1118^ and UAS-shi^ts^/+ in Figs. 5 and S5 were the same. In **(A-F, H-J),**the statistical significance was assessed by the comparison to UAS-shi^ts^/+ controls. In **(G-I, K-M),**the statistical significance was assessed by the comparison to w^1118^. *: p<0.01.

**Fig. S6.**
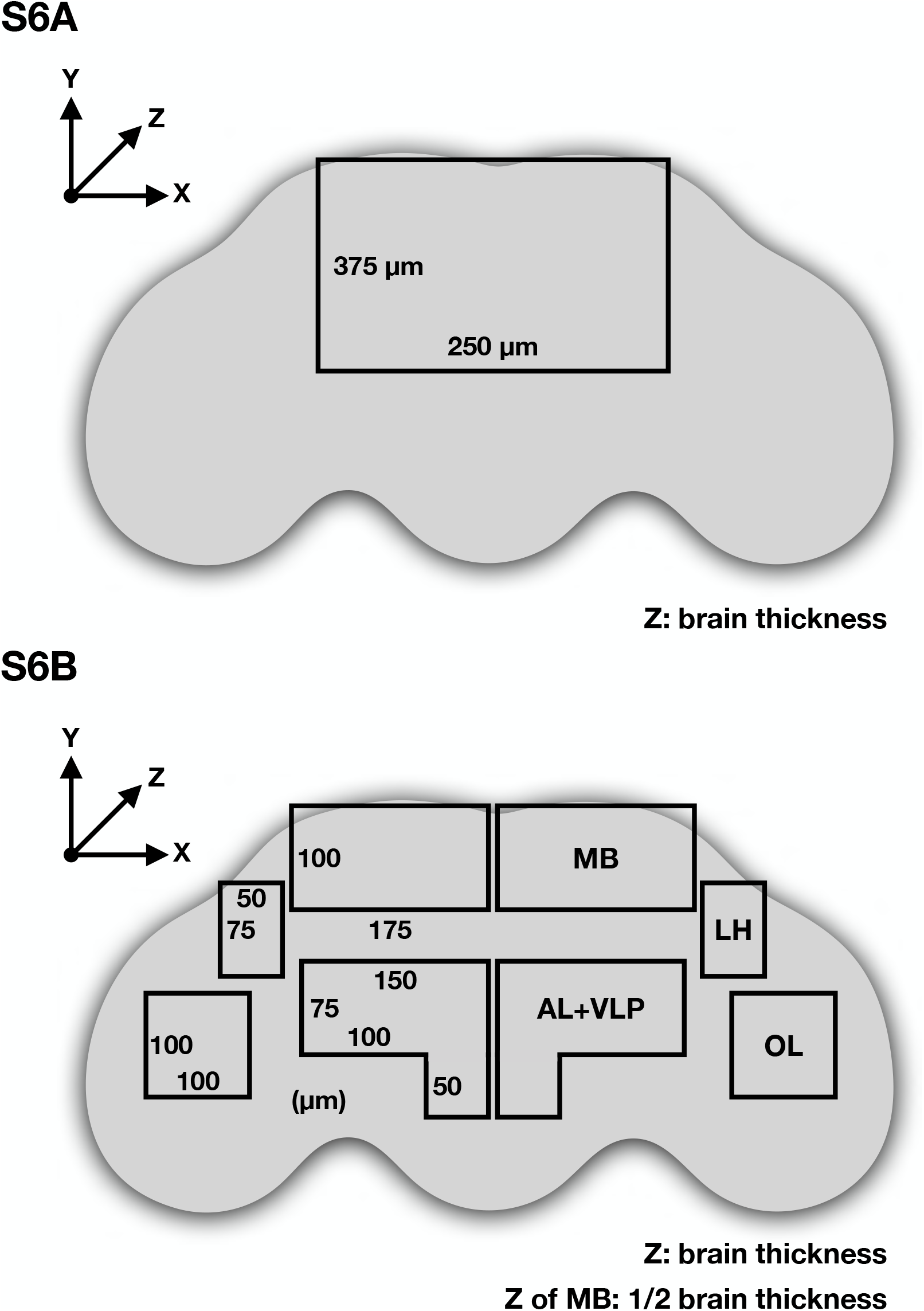
Schematic illustration of the brain regions analyzed. **(A)** and **(B)** The brain regions in the black outlines denoted the area where **(A)** 41Q-HA aggregation and **(B)** cleaved Caspase-3-positive cells were analyzed. **(A)** and **(B:**LH, AL+VLP, and OL) were analyzed with whole brain sections (1 μm/section). **(B:**MB) was analyzed with posterior 1/2 sections. MB, mushroom body; LH, lateral horn; AL+VLP, antenna lobe and ventrolateral protocerebrum; OL, optic lobe.

**Table S1. List of primer sequences used for generating hsp22-HA and miR-hsp22 lines**

**Mov. S1. DNIC-1 alleviates the polyQ-mediated motor dysfunction**

This video showed the DNIC-1-treated polyQ-expressing flies have better motor activities. The animals used were 23 day-old female flies. At this age, untreated polyQ-flies already displayed strong motor defects. Left, 0.5% DMSO (controls). Right, 500 μM DNIC-1. Genotype for the 41Q-HA-expressing flies: TubP-Gal80^ts^/+; nSyb-Gal4/UAS-41Q-HA.

## Notes

### Competing Interest Statement

The authors have declared no competing interest.

